# Panoptic vDISCO imaging reveals neuronal connectivity, remote trauma effects and meningeal vessels in intact transparent mice

**DOI:** 10.1101/374785

**Authors:** Ruiyao Cai, Chenchen Pan, Alireza Ghasemigharagoz, Mihail I. Todorov, Benjamin Förstera, Shan Zhao, Harsharan S. Bhatia, Leander Mrowka, Delphine Theodorou, Markus Rempfler, Anna Xavier, Benjamin T. Kress, Corinne Benakis, Arthur Liesz, Bjoern Menze, Martin Kerschensteiner, Maiken Nedergaard, Ali Ertürk

**Affiliations:** Institute for Stroke and Dementia Research, Klinikum der Universität München, Ludwig-Maximilians University Munich, Germany; Munich Cluster for System Neurology (SyNergy), 80336 Munich, Germany; Institute of Clinical Neuroimmunology, Klinikum der Universität München, Ludwig-Maximilians University Munich, Germany; Biomedical Center, Ludwig-Maximilians University Munich, Germany; Department of Computer Science & Institute for Advanced Study, Technical University of Munich, Munich, Germany; Center for Translational Neuromedicine, University of Rochester, NY 14642, USA; Center for Translational Neuromedicine, Faculties of Health and Medical Sciences, University of Copenhagen, 2200 Copenhagen, Denmark

## Abstract

Analysis of entire transparent rodent bodies could provide holistic information on biological systems in health and disease. However, it has been challenging to reliably image and quantify signal from endogenously expressed fluorescent proteins in large cleared mouse bodies due to the low signal contrast. Here, we devised a pressure driven, nanobody based whole-body immunolabeling technology to enhance the signal of fluorescent proteins by up to two orders of magnitude. This allowed us to image subcellular details in transparent mouse bodies through bones and highly autofluorescent tissues, and perform quantifications. We visualized for the first-time whole-body neuronal connectivity of an entire adult mouse and discovered that brain trauma induces degeneration of peripheral axons. We also imaged meningeal lymphatic vessels and immune cells through the intact skull and vertebra in naïve animals and trauma models. Thus, our new approach can provide an unbiased holistic view of biological events affecting the nervous system and the rest of the body.

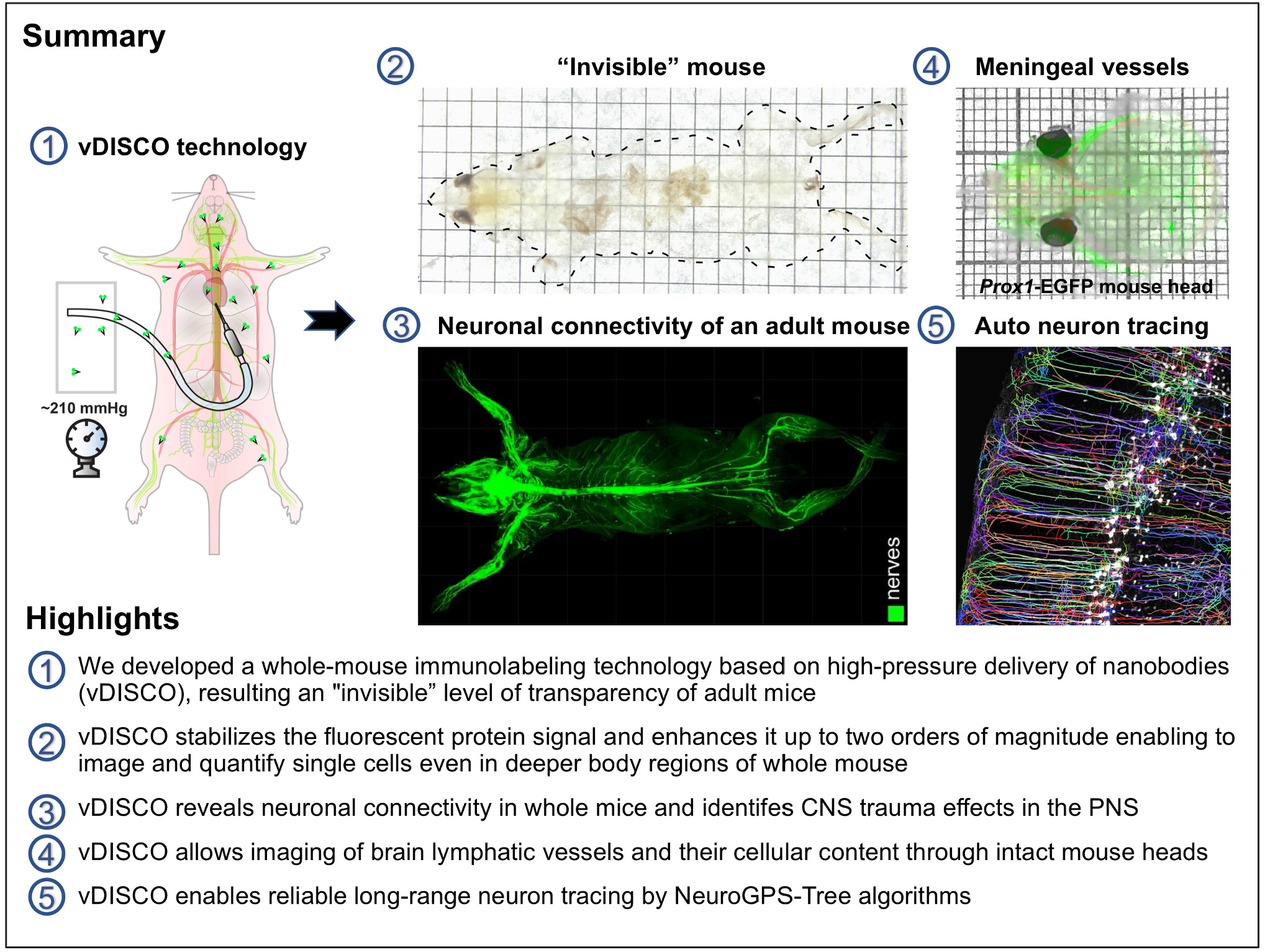

**Note: Manuscript videos are available at ‘Supplementary material’ section of BioRxiv and at the following link http://vdisco.isd-muc.de/**

## Introduction

Most diseases, even when arising in a specific lesion or locus, eventually come to affect the entire organism. Histological techniques developed in the last century have been the standard procedure for charting pathology, yet, a more complete understanding of biological mechanisms requires an unbiased exploration of the whole organism, not just selected tissue slices.

However, mammalian tissues are naturally opaque, hindering high-resolution imaging in any tissue deeper than a few hundred micrometers– a major reason why sectioning is needed for histological examination of target organs^1^. Recent innovations in tissue clearing technology now allow 3D histological examination of intact organs^2-15^. Tissue clearing is a chemical process aiming to match refractive indices throughout intact tissues, thus rendering them transparent and allowing deep tissue fluorescent microscopy. Most initial applications of tissue clearing have focused on clearing and imaging of endogenous fluorescent proteins expressed in the mouse brain. More recently, deep tissue immunolabeling methods have enhanced the quality of imaging for whole rodent organs as well as human embryos thanks to conjugation of bright fluorescent dyes with secondary antibodies^4, 16, 17^. A few studies have even rendered entire adult mouse bodies transparent^3, 18^, and allowed head-to-toe light-sheet microscopy imaging of intact adult mice^19^. All whole-body clearing and imaging methods up to date relies on transgene expression of fluorescent proteins such as EGFP, EYFP and mCherry^19, 20^. While these fluorescent proteins emit light in the visible spectrum, skeletal muscle and other bodily tissues have obstructive autofluorescence in this range^21^. In addition, fluorescent proteins are often less bright compared to most of the synthetic fluorophores and their signal intensity is further attenuated during the clearing and imaging procedures. Together, these bottlenecks hinder the reliable detection and quantification of subcellular details in centimeters-thick transparent mice, thereby needing imaging of dissected organs for quantifications, compromising the benefit of whole-body transparency.

Here, we developed a whole-body immunolabeling method to boost the signal of fluorescent proteins (named vDISCO). This technology enhances the fluorescent signals up to 118 times, and thereby allows panoptic imaging and quantification of subcellular details in transparent mice including changes in neuronal connectivity at the neuromuscular junctions upon brain trauma.

## Results

### vDISCO principles and signal enhancement

We reasoned that enhancing the signal of fluorescent proteins via whole-body immunolabeling with brighter and more stable fluorescent dyes could provide a much higher contrast (signal-to-background ratio = SBR) in cleared mice. Furthermore, using fluorescent dyes with emission peaks in the far-red range could help to overcome tissue autofluorescence, thus affording reliable detection of subcellular details in all tissues^4, 21^ (**Supplementary Fig. 1**). We reasoned that nanobodies are particularly suited to achieve a thorough immunolabeling throughout adult mice because of their small molecular weight (12-15 kDa) compared to that of conventional antibodies (~150 kDa)^3, 22^. For whole-body labeling, we delivered anti-XFP nanobodies conjugated to bright Atto dyes (called nanoboosters) using a pressure driven peristaltic pump to enhance their penetration deep into tissues.

We first tested the signal quality of nanobody labeling in dissected mouse brains coming from *Thy1*-GFPM mice, in which a subset of neurons express EGFP^23^. Following the nanoboosting, the tissues were cleared using organic solvents^6, 7^. We found that nanoboosting enhanced the signal quality one to two order of magnitude compared to direct imaging of fluorescent proteins (**Fig. 1a-k**). In the nanoboosted samples, fine details of neurons were evident even in the regions with fewer neurons labelled in *Thy1*-GFPM mice such as the midbrain (**Fig. 1c,d** vs. **Fig. 1g,h**). We obtained a similar signal quality increase in the cerebellum of *Thy1*-GFPM brains (**Fig. 1i** vs. **Fig. 1j**), which is notoriously difficult to clear due to the high lipid content.

**Figure 1.**
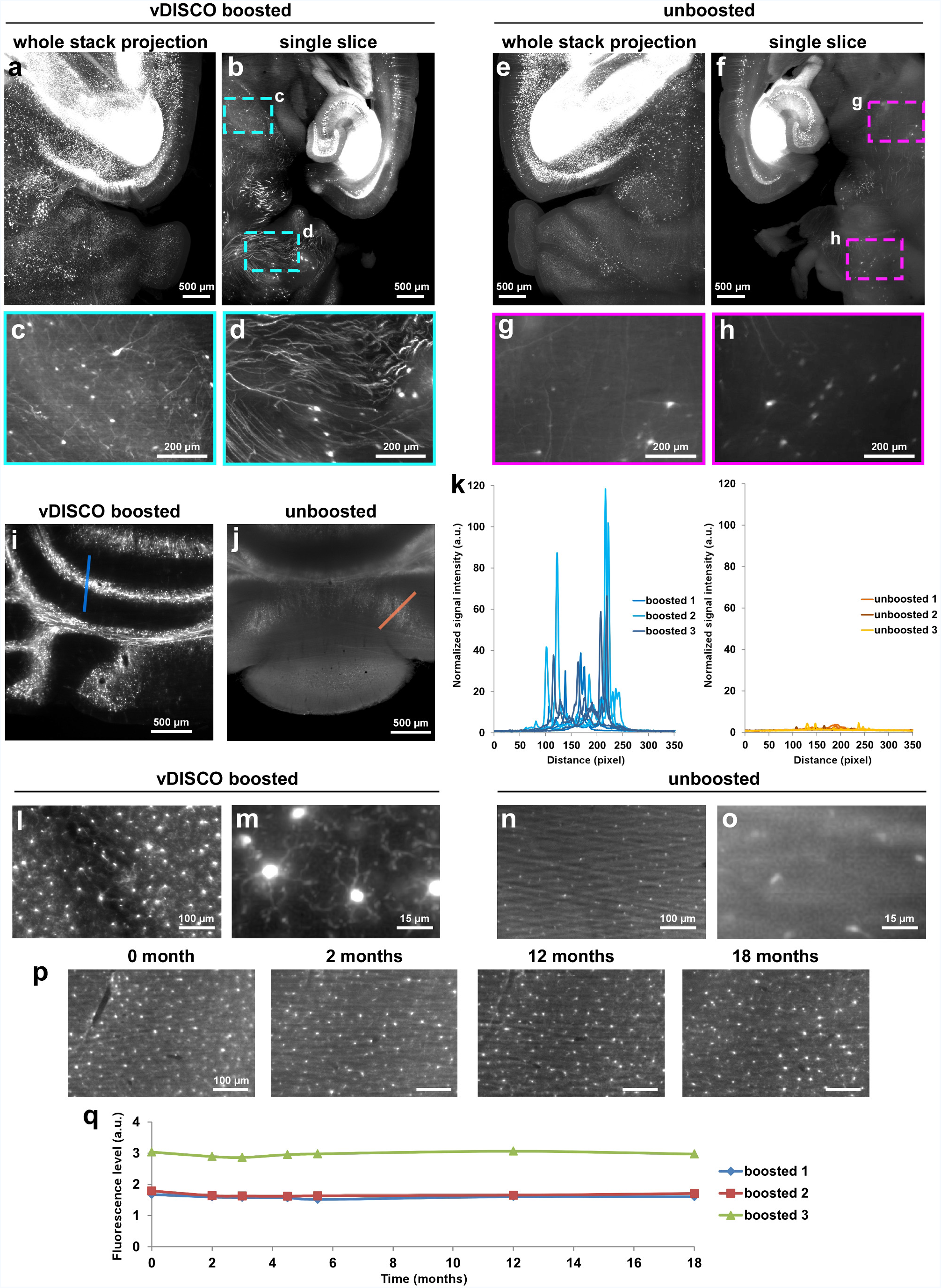
Enhanced signal and details in Thy1-GFPM and CX3CR1^GFP/+^ mouse and permanent preservation of the fluorescence signal with vDISCO. Signal quality comparison between vDISCO boosted (**a**-**d**) and unboosted (**e**-**h**) half brain samples coming from the same Thy1-GFPM mouse. To achieve the best comparison between the two procedures, we divided the mouse brains in two halves for light-sheet microscopy imaging (one hemisphere boosted and imaged in far-red channel, the other hemisphere unboosted and imaged in the green channel for endogenous EGFP). **b** and **f** are single images from the maximum intensity projections in a and e, respectively. The boosted hemisphere showed highly distinguishable cellular details (**c**, **d**) such as axonal projections not visible in the unboosted hemisphere (**g**, **h**) especially in the regions with dim EGFP labeling in Thy1-GFPM mice, such as mid brain and cerebellum. (**i**, **j**) Comparison of signal quality in cerebellum from the boosted (**i**) vs. unboosted (**j**) samples. (**k**) Plots of signal intensity profiles from boosted samples (left) vs. unboosted samples (right) along the blue and orange lines in panels **i** and **j**, respectively. n=3 brains for each group (2-6 months old Thy1-GFPM mice). (**l**-**o**) 4x and 25x magnification light-sheet microscopy images of the microglia from CX3CR1^GFP/+^ in boosted (**l**, **m**) vs. unboosted samples (**n**, **o**) showing the fine details of microglia ramifications obtained with vDISCO boosting. (**p**) Light-sheet microscopy images of a CX3CR1^GFP/+^ mouse brain at 0, 2, 12 and 18 months after boosting, showing the preservation of the fluorescence signal over 18 months. (**q**) Fluorescence level quantifications in CX3CR1^GFP/+^ brains after boosting at different time points post-clearing (n=3 animals).

A major aim of tissue clearing approaches is to perform automated quantifications in large imaging datasets in an unbiased and timely way. Towards this goal, we used the NeuroGPS-Tree algorithm, a robust automated neuron-tracing tool that was recently developed for tracing cortex regions obtained by high-resolution confocal microscopy^24^. We found that, virtually all of the neuronal cell bodies and neurites were detected and linked to each other as complete neurons upon nanoboosting (**Supplementary Fig. 2**). In contrast, in unboosted samples, many fine extensions of neurons were not identified or not connected to neuronal trees (**Supplementary Fig. 2**). Nanoboosting allowed imaging of not only neuronal details, but also smaller individual cells such as immune cells (**Supplementary Fig. 3**). Compared to unboosted samples, we could resolve fine details of microglia cells in intact transparent brains of CX3CR1^GFP/+^ mice^25^ using light-sheet microscopy (**Fig. 1l-o**). Nanoboosting also enabled automated quantification of CX3CR1 GFP+ cells: using ClearMap algorithms^26^ we found approximately 2.59 million microglia cells in the adult mouse brain. We also automatically quantified microglia in all brain regions annotated by the Allen brain atlas. Thereby we found for example ~150,000 microglia in the hippocampus and ~50,000 in the thalamus of adult mice (**Supplementary Fig. 4**). Furthermore, we found no significant decrease of signal quality suggesting that nanoboosting stabilizes the fluorescent signal (**Fig. 1p,q**). Thus, owing to the enormous enhancement and stabilization of signal with nanobodies, we could use available computational tools to automatically trace neurons and count cell numbers in even low resolution light-sheet microscopy imaging datasets.

### vDISCO allows panoptic imaging of intact adult mouse bodies

To understand how the organisms function at the systems biology level, it is critical to obtain a subcellular resolution view not only on single organs but also on intact organisms. Towards this goal, we established an approach to achieve nanoboosting in the entire mouse body (**Fig. 2a**, **Supplementary Fig. 5**). We used a pressure delivery of a permeabilization solution containing Triton X-100, methyl-β-cyclodextrin (to extract the cholesterol from membranes), and trans-1-acetyl-4-hydroxy-L-proline (to loosen the collagen network)^27^. To further reduce the background caused by the residual blood and to decalcify the bones, we treated whole mouse bodies with aminoalcohols^18^ and EDTA^20, 28^, before the whole-body immunolabeling step. In addition to the specific boosted signal, we also visualized other major tissues in the transparent body: muscles by their autofluorescence at the blue-green spectra, bones and internal organs by propidium iodide (PI) labeling (**Fig. 2b-c, Supplementary Fig. 6a-d**). Usage of organic solvents inducing shrinkage^19^ allowed us to perform a head-to-toe panoptic imaging of a whole mouse by light-sheet microscopy.

**Figure 2.**
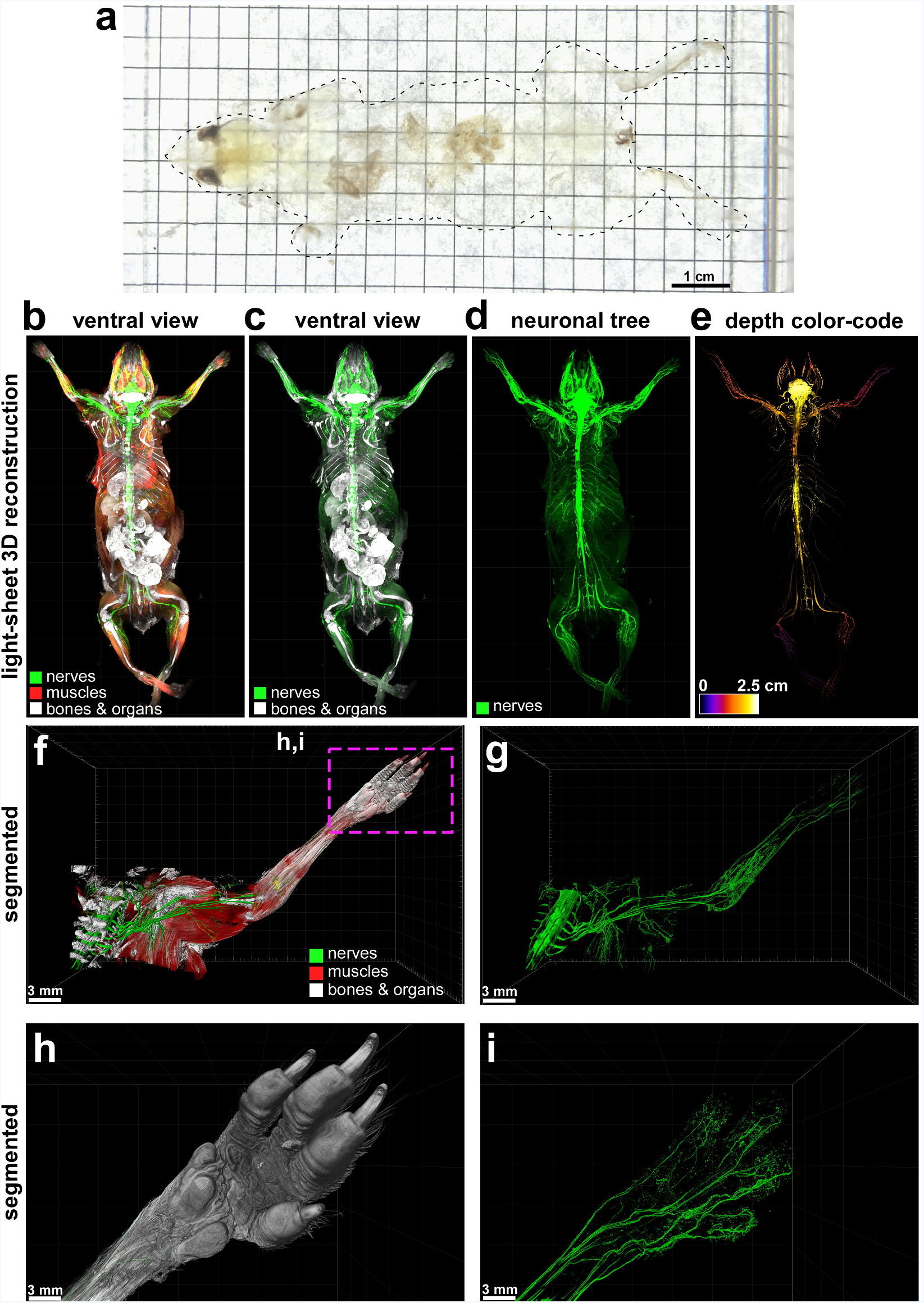
Panoptic imaging of intact Thy1-GFPM mouse. (**a**) An example of “invisible mouse” generated by vDISCO. (**b**-**e**) Whole body 3D reconstructions from light-sheet microscopy imaging of a Thy1-GFPM animal. vDISCO boosted EGFP+ neuronal structures are shown in green, bones and internal organs that are prominent with PI labeling in white, and the muscles visualized by autofluorescence background imaging are in red (**a**-**c**). The depth color-coding shows the neuronal projections at different z-levels in 2.5 cm thick whole mouse body (**d**). (**f**, **g**) High resolution 3D reconstruction views of the left torso and forelimb from the same animal in a-d. Details of innervation throughout muscles and bones are evident. (**h**, **i**) The surface reconstruction of the paw (**h**) and its nerves (**i**) from the marked region in e. See also Video 1-3.

Being able to image subcellular details of neurons through intact bones and highly autofluorescent muscles in whole mouse body with vDISCO panoptic imaging, we constructed the first whole-body neuronal connectivity map of a *Thy1*-GFPM transgenic mouse (**Fig. 2 d,e Supplementary Fig. 6, Video 1**). We noticed that in the peripheral nervous system (PNS) mainly axons innervating neuromuscular junctions were labelled in this mouse line. We also observed fluorescent labeling of the internal organs in rare cases such as kidneys of *Thy1*-YFPH mice, which was not reported before^23^ (**Supplementary Fig. 7**). Owing to the great increase in SBR by vDISCO, we could readily visualize details of axonal extensions from spinal cord through intact vertebra, until their terminals into the muscles and toes (**Fig. 2f-i, Videos 2,3**). Panoptic imaging of individual neuronal connections through intact bones can now provide more details on neuroanatomy. For example, how axons coming from consecutive roots enter the spinal cord i.e., in an overlapping or non-overlapping manner has been unclear^29^. Using our technology, we observed that in mice, the axonal bundles coming from different roots enter the spinal cord at non-overlapping territories (**Supplementary Fig. 8**). Thus, vDISCO approach provides a holistic view of the intact mouse, which should lead to novel discoveries on how interconnected organ systems function in health and what happens during their perturbation in disease.

### Degeneration of peripheral nerve terminals after brain injury

Traumatic brain injury (TBI) is a major cause of death and disability and currently there is no disease modifying treatment available. It often leads to chronic focal and global neurological impairments, such as dementia, epilepsy and progressive motor decline^30^. However, it is unclear whether and where neuronal connectivity is altered in remote body regions upon acute brain injuries. Here, we used panoptic imaging to examine neuronal changes throughout the body more than a month after brain injury in comparison to control animals. As mainly axonal extensions innervating muscles are labelled in *Thy1*-GFPM mice, we primarily focused on these nerve terminals. Our quantifications on intact cleared mouse bodies demonstrated that the complexity of nerve terminals at the neuromuscular junctions was reduced after TBI, especially in the upper torso compared to unlesioned control mice (**Fig. 3a-d**). We observed that nerve endings were largely reduced in size, with fewer axonal ramifications left, implying partial degeneration of these axon terminals (**Fig. 3e,f**). These data show that light-sheet microscopy imaging data of transparent mouse bodies by vDISCO can be quantified to obtain previously unknown biological information.

**Figure 3.**
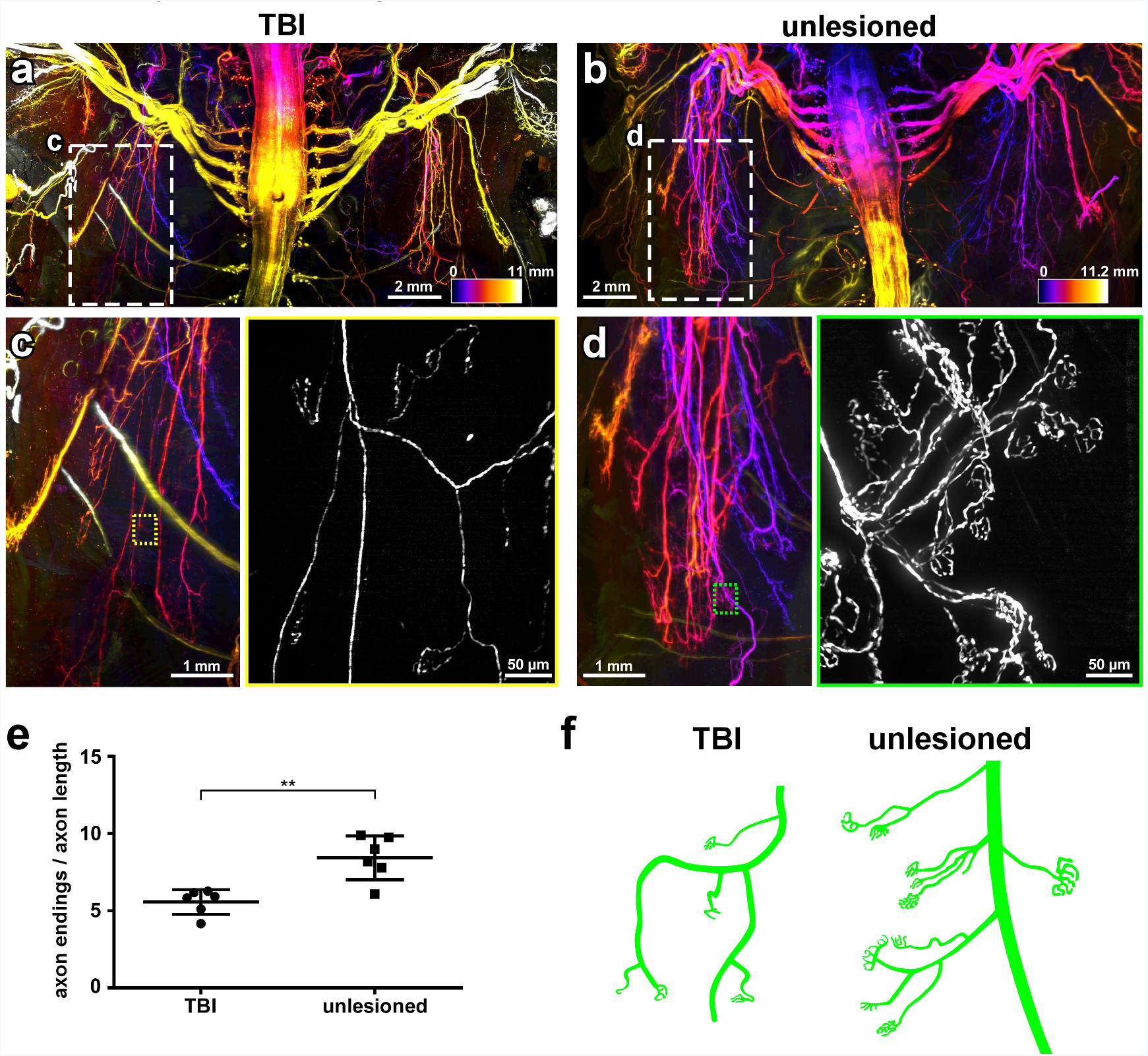
Panoptic imaging of peripheral nerve degeneration induced by traumatic brain injury (TBI) Light-sheet microscopy images of the torso from TBI-induced (**a**) vs. unlesioned control (**b**) Thy1-GFPM mice. The color-code indicates the z-depth of anatomical regions as given in the scale bars. (**c**, **d**) High magnification views of marked regions in a and b, showing the left toracic peripheral nerve projections in the TBI (**c**) vs. control animal (**d**). The colored rectangles show the high magnification images of the marked regions in c and d, respectively, demostrating fewer intact peripheral nerve endings in TBI mice compared to controls. (**e**) Quantification of axonal projection complexity expressed as number of peripheral nerve endings over length of axonal ramifications in TBI vs. control mice (mean ± s.d.; n=6 animals per group; statistical significance (**p < 0.01) was assessed by two tailed t-test). (**f**) Representative illustration showing the peripheral nerve ending morphology in TBI vs. control mice.

### Imaging meningeal vessels of the central nervous system (CNS)

The lymphatic system, which connects various lymphatic organs in the body, is crucial for immune responses. Until recently, the brain was considered to be devoid of any lymphatic vessels. In the recent years, however, existence of brain lymphatic vessels was discovered, which calls for a re-evaluation of immune cell trafficking routes between the CNS and the rest of the body^31-33^. As these lymphatic vessels are located between the brain and skull, their connections are largely destroyed when the brain is harvested for standard histology.

We utilized panoptic imaging to overcome this hurdle and image details of meningeal vessels in intact transparent mice. We first used *Prox1*-EGFP reporter mice, in which lymphatic endothelial cells express EGFP^34^. We readily observed previously described brain lymphatic structures in complete, particularly along the sagittal sinus and pterygopalatine artery (**Fig. 4a-d**). Next, we injected a cerebral spinal fluid (CSF) tracer (fluorophore-conjugated ovalbumin) into the cisterna magna in *VEGFR3*-YFP mice, another commonly used reporter line for meningeal vessels. We could clearly visualize vascular connections between the brain and skull, which were labelled with CSF tracer (**Fig. 4e-h**). Next, we imaged multiple subtypes of immune cells within and outside the meningeal vessels using the CX3CR1^GFP/+^ x CCR2^RFP/+^ mice by multiplexing with two different nanoboosters (anti-GFP conjugated to Atto-647N and anti-RFP conjugated to Atto-594N). The CX3CR1 GFP+ microglia cells in the brain parenchyma (**Fig. 4i,j**, cyan arrowheads) and CCR2 RFP+ peripheral immune cells in the meningeal vessels were clearly identifiable (**Fig. 4i,j**, white arrowheads and **Video 4**). As expected, we observed CX3CR1 GFP+ immune cells also in meningeal vessels, which likely represent the peripheral macrophages and monocytes (**Fig. 4j**, yellow arrowheads). Thus, vDISCO panoptic imaging of transparent mice is a powerful tool to study the intact anatomy of meningeal vessels in their native environment.

**Figure 4.**
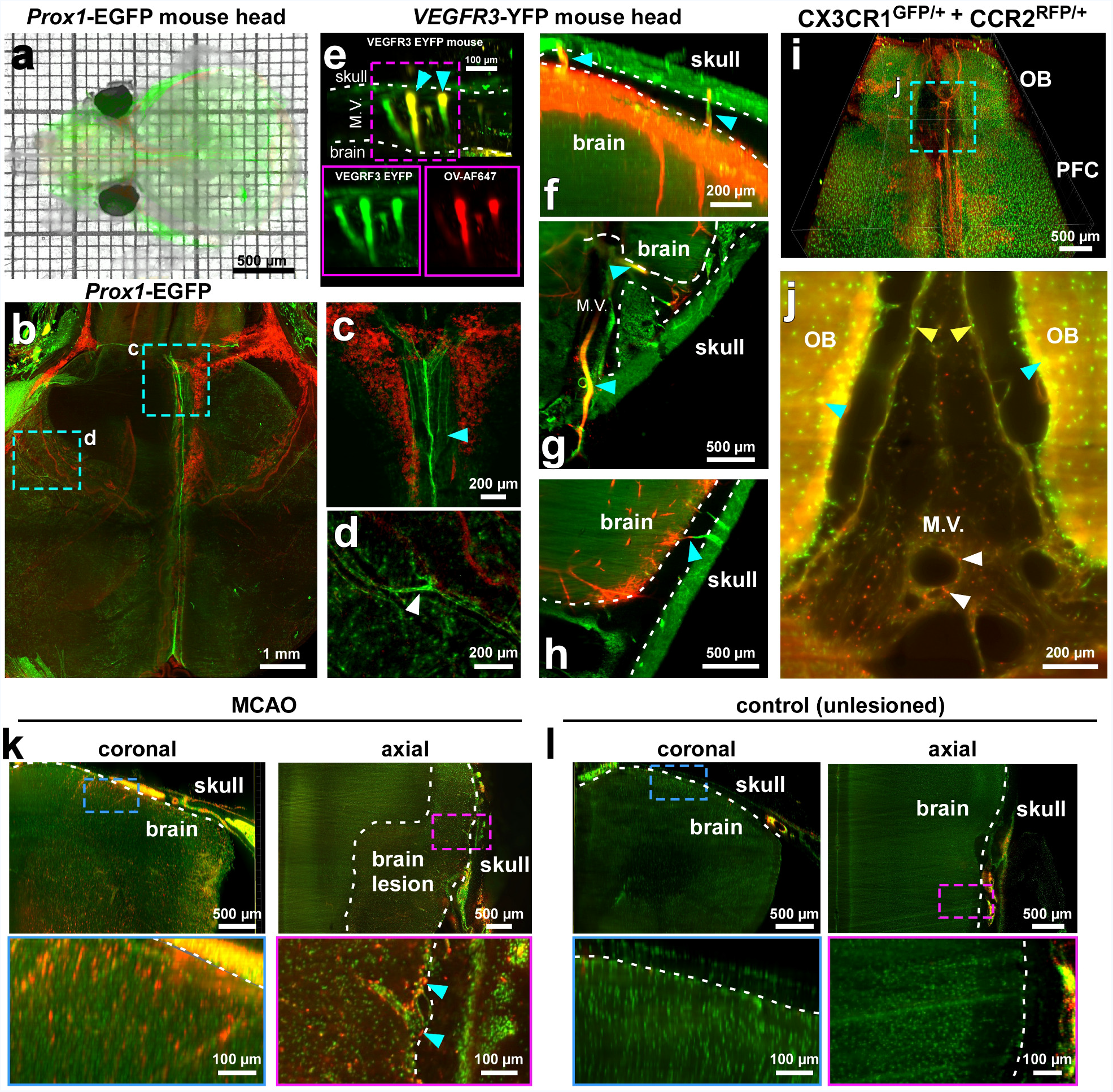
Panoptic imaging of meningeal vessels and immune cells through transparent skull. (**a**) A transparent Prox1-EGFP mouse head showing the labeled vessels underneath the skull. (**b**-**d**) The Prox1-EGFP mouse head showing the brain lymphatic vessels along the sagittal sinus and pterygopalatine artery (cyan and white arrowheads in **c** and **d**, respectively). Bone structures become prominent with PI labeling (red). VEGFR3-YFP mouse heads in sagittal (**e**, **f**) and axial views (**g**, **h**) showing the meningeal vessels (M.V., green) along with the CSF tracer (Ovalbumin-Alexa647, red) labelling mainly lymphatic vessels (cyan arrowheads). (**i**) 3D visualization of prefrontal cortex (PFC) and olfactory bulb (OB) in a CX3CR1^GFP/+^ (green) and CCR2^RFP/+^ (red) double transgenic mouse. (**j**) High-magnification image of marked region in i showing CCR2 RFP+ cells (white arrowheads) and CX3CR1 GFP+ cells (yellow arrowheads) in meningeal vessels, and the CX3CR1 GFP+ cells in the brain parenchyma (cyan arrowheads). (**k**, **l**) LysM-EGFP transgenic mice with MCAO stroke model vs. unlesioned control showing the infiltration of immune cells in MCAO. LysM GFP+ cells are shown in red and nucleus labeling by PI in green. Immune cells in the meningeal vessels of the injured mice were observed (cyan arrowheads). All images were obtained by light-sheet microscopy. See also Video 4.

It remains unclear, how the immune cells within meningeal vessels contribute to the pathology of acute brain injuries (or other neurological diseases), although they may be in an advantageous position to invade the brain. To start addressing this query, we used the middle cerebral artery occlusion (MCAO) model of stroke, which was performed on *LysM*-EGFP mice, a transgenic mouse line expressing EGFP in neutrophils, monocytes and macrophages but not in microglial cells^35^. The mice with MCAO showed an invasion of LysM GFP+ cells into the brain parenchyma especially in the peri-infarct region compared to sham controls (**Fig. 4k,l**). LysM GFP+ cells also increased in the meningeal structures of MCAO mice (**Fig. 4k** cyan arrowheads) suggesting that the meninges might be playing a role as entry and/or exit route of the brain in addition to the disrupted blood brain-barrier and the choroid plexus^36^. Finally, we explored the inflammatory reaction after spinal cord injury (SCI) using panoptic imaging. To this end, we induced a SCI in *CD68*-EGFP transgenic mice expressing EGFP in monocytes and macrophages^37^. Upon SCI, we observed an influx of CD68 GFP+ cells at the spinal lesion site as well as a rather widespread increase in CD68 GFP+ cells around the site of the spinal cord trauma including in the adjacent muscles, spinal cord roots and meningeal vessels (**Supplementary Fig. 9, Video 5**) suggesting that a broader view on inflammation in SCI pathology is needed.

## Discussion

Imaging of transparent adult mouse bodies from head to toe holds the promise of providing an unbiased and highly resolved view of entire organ systems in health and disease. Here, we developed a new whole-body nanobody labeling method in conjunction with whole-body tissue clearing, enabling a reliable visualization and quantification of subcellular details throughout the centimeters-thick tissues of intact mouse bodies. This new panoptic imaging technology is straightforward in application, and suitable for systemic analysis of a vast range of biomedical inquires, as demonstrated here in the mapping of the neuronal connectome, and investigation of meningeal vessels and immune cell infiltrations through intact bones in acute CNS trauma models.

The panoptic imaging of nervous and immune systems in intact mouse bodies was achieved owing to the enhancement of signal quality by one to two orders of magnitude via whole-body immunolabeling with nanobodies conjugated to bright dyes in far-red spectra. Therefore, autofluorescence in shorter wavelength spectra, in which traditional fluorescent proteins such as GFP are excited and imaged, is avoided. Here, we primarily used Atto dyes at 647 nm, and additional Atto 594 nm for multiplexing 2 different cell types. In the future, usage of nanobodies conjugated to dyes that emit in the near infrared range^38^ can further facilitate multiplexed detection of more than 2 targets. While the number of commercially available nanobodies is still limited, the nanoboosters used in this study can stain a broad selection of 21 different fluorescent proteins including EGFP, YFP, Venus, mCherry, and mRFP. Here, we used transgenic mice endogenously expressing fluorescent proteins, however, the labeling can also be achieved by rabies virus retrograde tracing, systematic AVV injections^39^ or transplantation of genetically engineered cells (such as stem cells or adoptive transfer of immune cells). Another key advantage of vDISCO panoptic imaging is that upon staining with Atto dyes through nanoboosting the signal becomes permanent, permitting long-term imaging sessions and also future re-analysis of the same sample as needed. It is notable that, while we developed vDISCO for panoptic imaging of the whole mouse body, it is readily applicable for individual organs, using a simplified protocol to drastically increase and stabilize the signal contrast, as we show for mouse brains imaged more than a year after clearing (**Video 6**).

A current limitation is the absence of light-sheet microscope systems that could image the entire body of a mouse in one session without any displacement and turning of the sample. Construction of such new light-sheet microscopes should further decrease the time needed for data acquisition and simplify data processing for 3D reconstruction. In fact, as shown here, optics and light sources of current commercial light-sheet microscopes are sufficient to image intact mouse bodies at subcellular resolution, albeit the whole body has to be manually moved in the imaging chamber to achieve the scanning of different body regions. In addition, development of algorithms to automatically identify and quantify cellular changes on the whole organism scale will be important to further scale up this technology in the future.

Using panoptic imaging, we constructed a neuronal connectivity map for the *Thy1*-GFPM transgenic mouse, showing subcellular details of long-range neuronal connections from CNS to the most distal body regions. Here, we visualized neuronal details in *Thy1*-GFPM mice, where 5-10% of CNS neurons and PNS neurons innervating skeletal muscles are labelled. In the future, panoptic imaging of other transgenic lines labeling a greater and more diverse subset of neurons will allow the exploration of other biological systems such as the autonomic nerve innervation of internal organs.

In recent years, researchers have been providing more evidences on post TBI changes that seem to form the basis of chronic complications, such as epilepsy, neuropsychiatric disorders, dementia and progressive motor decline among others^30^. However, the effects of a localized brain lesion on the rest of the body have been poorly understood mainly due to the technical challenges in studying long-range neuronal connectivity. Because vDISCO fluorescent signal boosting allows both imaging and quantifications on light-sheet microscopy images of transparent mouse bodies, we identified TBI-induced changes at the peripheral axonal projections innervating the skeletal musculature of the torso. These data are in line with our recent results demonstrating functional deficits in fine motor movements in mice upon TBI in the same experimental model^40^. Here, we also demonstrated that meningeal vessels extending between the cranium and CNS tissue and their cellular content can readily be imaged by vDISCO panoptic imaging in naïve animals and acute CNS injury models (stroke and spinal cord injury). As neuroinflammation is a major determinant of neuronal function and survival following CNS insults of various aetiologies, our technology should accelerate the study of brain lymphatic vessels in various CNS pathologies.

Thus, vDISCO enhancement and stabilization of fluorescent proteins combined with signal acquisition in far-red spectra should facilitate discovery of previously inaccessible biological information. For example, imaging and quantification of neuronal connectivity and immune cell populations in the entire mouse body should contribute to a more comprehensive understanding of the degenerative and inflammatory pathways in disease states such as dementia and neuropsychiatric disorders.

## Acknowledgements

This work was supported by the Vascular Dementia Research Foundation, Synergy Excellence Cluster Munich (SyNergy), ERA-Net Neuron (01EW1501A, A.E.), NIH (A.E. and M.N.), the Novo Nordisk Foundation (M.N.), the Howard Hughes (B.T.K.), and the Lundbeck Foundation (to A.X. and M.N.). We thank Antonia Weingart for illustrations, Farida Hellal for technical advice and critical reading of the manuscript, and Arnaldo Parra Damas, Francesca Quacquarelli, and Giuseppe Locatelli for help during initial optimization. A.E, C.P., R.C., A.L. and M.I.T. are members of the Graduate School of Systemic Neurosciences (GSN) at the Ludwig Maximilians University of Munich.

## Author contributions

A.E. initiated and led all aspects of the project. R.C. and C.P. developed the method and conducted most of the experiments. R.C. and A.G. analyzed data. M.I.T. stitched and analyzed the whole mouse body scans. B.F. and S.Z. helped to optimize the protocols. H.S.B., L.M., M.R. and B.M., analyzed data. D.T. and M.K. contributed spinal cord injury experiments; C.B. and A.L. MCAO experiments; and A.X., B.K., and M.K., CM injection experiments. A.E. wrote the paper. All the authors edited the manuscript.

## MATERIALS AND METHODS

### Animals

We used the following animals in the study: mixed gender CX3CR1^GFP/+^ (B6.129P-Cx3cr1tm1Litt/J, Jackson Laboratory strain code: 005582), *Thy1*-GFPM, *Thy1*-YFPH^23^, *Prox1*-EGFP (Tg(Prox1-EGFP)KY221Gsat/Mmucd, MMRRC strain code: 031006-UCD), *VEGFR3*-YFP, CX3CR1^GFP/+^ x CCR2^RFP/+^ (B6.129(Cg)-Ccr2tm2.1Ifc/J, Jackson Laboratory strain code: 017586), *LysM*-EGFP (Lyz2tm1.1^Graf^, MGI: 2654931), *CD68*-EGFP (C57BL/6-Tg(CD68-EGFP)1Drg/j), Jackson Laboratory strain code: 026827). The animals were housed under a 12/12 hours light/dark cycle. The animal experiments were conducted according to institutional guidelines: Klinikum der Universität München / Ludwig Maximilian University of Munich and after approval of the Ethical Review Board of the Government of Upper Bavaria (Regierung von Oberbayern, Munich, Germany) and the Animal Experiments Council under the Danish Ministry of Environment and Food (2015-15-0201-00535) and in accordance with the European directive 2010/63/EU for animal research. All data are reported according to the ARRIVE criteria. Sample sizes were chosen based on prior experience with similar models. Sample sizes are specified in figure legends. Within each strain, animals were randomly selected.

### Traumatic brain injury

Traumatic brain injury was performed using a Controlled Cortical Impact (CCI) device (Leica Benchmark Stereotaxic Impactor, 39463923). 30 minutes before surgery, we administered Carprofen (4 mg/kg) and Buprenorphin (0.05 mg/kg) to the animals via subcutaneous injection. Then, anaesthesia was induced in animals with 4% isoflurane in N_2_O/O_2_ (70%/30%) mixture and afterwards maintained with 1.5% isoflurane in the same mixture for the whole surgery. As soon as the animals did not show any pedal reflex, they were placed in the associated stereotaxic apparatus and their body temperature was kept at 37°C using a heating pad for the whole surgery procedure. Next, the scalp of the animals was shaved, aseptically prepared by wiping with Octenisept (Schülke, 22580-A) as disinfectant and the skin of the scalp was incised longitudinally between the occiput and forehead. We identified the target area of the injury, which was the right somatosensory cortex, using the stereotaxic frame (bregma coordinates: 2-mm posterior, 5-mm right lateral). The injury was then triggered via CCI machine using the following parameters: impact speed: 6.9 m/s; impact duration: 400 ms and impact depth: 2 mm. With these parameters, the injury resulted severe with cracks of the skull, bleeding and exposed brain tissue. After the impact, the skin was sutured with metallic wound closure clips (VWR, 203-1000) and the animals were kept at 31°C in a recovery chamber (Mediheat, 34-0516) for at least 30 minutes until they recovered from the anesthesia. In the following days, animals were subcutaneously injected with Carprofen (4 mg/kg) once every day for 4 days and sacrificed at > 1 month post injury by transcardial perfusion according to the ‘perfusion and tissue preparation’ section below.

### MCAO model

Experimental stroke was induced using the intraluminal filament model of middle cerebral artery (MCA) occlusion for transient, focal brain ischemia. Mice were anesthetized with isoflurane delivered in a mixture of 30% O_2_ and 70% N_2_O. A heat-blunted nylon suture (6/0) was inserted into the external carotid artery and advanced until it obstructed the MCA together with the ligation of the common carotid artery for 30 min. Regional cerebral blood flow (CBF, bregma coordinates: 2-mm posterior, 5-mm lateral) was continuously recorded by transcranial laser Doppler flowmetry from the induction of ischemia until 10 min after reperfusion. Following fMCAO, mice were placed in temperature-controlled recovery cages for 2 h to prevent post-surgery hypothermia. For the survival period (3 days), the mice were kept in their home cage with facilitated access to water and food. Sham-operated mice received the same surgical procedure without insertion of the filament. Body temperature was maintained throughout surgery using a feedback-controlled heating pad and kept constant (37.0 ± 0.5 °C). Exclusion criteria were as follows: insufficient MCA occlusion (a reduction in blood flow to 15% of the baseline value) and blood flow recovery >80% within 10 min of reperfusion. Mice were sacrificed at 3 days post injury by transcardial perfusion according to the ‘perfusion and tissue preparation’ section below.

### CM injection for meningeal vessel labeling

For the cisterna magna injections, mice (*VEGFR3*-YFP; 6 months old) were anesthetized with a mixture of ketamine and xylazine (100 mg/kg; 10 mg/kg, respectively) via intraperitoneal (i.p.) injection. Upon toe pinch reflexes ceased, mice were fixed in a stereotaxic frame by the zygomatic arch, with the head slightly tilted to form an angle of 120° in relation to the body. The head and neck regions were shaved to expose the neck muscles, which were bluntly dissected to expose the cisterna magna (CM). Cannulas composed of a dental needle (SOPIRA^®^ Carpule 30G 0.3 × 12mm; Kulzer; AA001) and polyethylene tubing (0.024” OD x 0.011” ID; Scandidact; PE10-CL-500) were used to perform the CM injections. A cannula filled with cerebrospinal fluid (CSF) tracer (2% ovalbumin 45 kDa – Alexa Fluor 647 conjugated; Thermo Fisher Scientific, O34784, diluted in artificial CSF: 126 mM NaCl, 2.5 mM KCl, 1.25 mM NaH_2_PO_4_, 2 mM Mg_2_SO_4_, 2 mM CaCl_2_, 10 mM glucose, 26 mM NaHCO_3_; pH 7.4 when gassed with 95% O_2_ and 5% CO_2_) was inserted into the CM. With the aid of an injection pump (LEGATO 130 Syringe pump, KD Scientific, 788130), 10 μl of CSF tracer was injected into the CM at a rate of 1 μl/min. At the end of the injection, CSF tracer was allowed to circulate in the subarachnoid and paravascular spaces for 1 hour. Mice were then transcardially perfused according to the ‘perfusion and tissue preparation’ section below.

### Spinal cord injury model

Mice were deeply anaesthetized by i.p. injection of a combination of midazolam (5 mg/kg body weight), medetomidine (0.5 mg/kg) and fentanyl (0.05 mg/kg). The mid-thoracic spinal cord of anaesthetized *CD68*-EGFP mice was surgically exposed by a dorsal laminectomy as previously described^41^. A hemisection of the spinal cord was performed using fine-tip surgical scissors (F.S.T 15000-08 spring scissor 2,5 mm cutting edge). Muscle tissue and skin were then sutured with a surgical thread (Ethilon suture 6-0, 667H) and animals were allowed to recover on a heating pad. After 48 hours, mice were transcardially perfused according to the ‘perfusion and tissue preparation’ section below.

### Perfusion and tissue preparation

Mice were deeply anesthetized using a combination of midazolam, medetomidine and fentanyl (MMF) (1ml/100g of body mass for mice; i.p.). As soon as the animals did not show any pedal reflex, they were intracardially perfused with heparinized 0.1 M PBS (10 U/ml of Heparin, Ratiopharm; 100-125 mmHg pressure using a Leica Perfusion One system) for 5-10 minutes at room temperature until the blood was washed out, followed by 4% paraformaldehyde (PFA) in 0.1 M PBS (pH 7.4) (Morphisto, 11762.01000) for 10-20 minutes Next, skin, optionally eyes, premaxilla and maxilla bones were carefully removed and the palate of the animal was opened (without damaging the tissue beneath), and the feces were gently washed out from the intestine with 0.1 M PBS through small cuts using a syringe. Then, the bodies were post-fixed in 4% PFA for 1 day at 4 °C and later washed with 0.1 M PBS for 10 minutes 3 times at room temperature. The whole-body nanoboosting procedure was started immediately or whole mouse bodies were stored in PBS at 4 °C for up to 4 weeks or in PBS containing 0.05% sodium azide (Sigma, 71290) for up to 6 months.

For the quantification of fluorescence signal, CX3CR1^GFP/+^ and *Thy1*-GFPM mice were perfused with PBS and PFA as described above. Subsequently, brains from these animals were dissected out and post-fixed in 4% PFA overnight at 4°C, washed with 0.1 M PBS for 10 minutes 3 times at room temperature and kept in PBS plus 0.05% sodium azide up to 3 weeks.

### Clearing of unboosted samples

For the quantification of fluorescence signal after clearing, without boosting, we followed the uDISCO passive clearing protocol as described in Pan *et al*., 2016^19^. In brief, dissected brains were placed in 5 ml tubes (Eppendorf, 0030 119.401) and covered with 4.5 ml of clearing solution. All incubation steps were performed in a fume hood with gentle shaking or rotation, with the samples covered with aluminum foil to keep them in dark. To clear the samples, we incubated them in a gradient of *tert*-butanol (Sigma, 360538): 30 Vol%, 50 Vol%, 70 Vol%, 80 Vol%, 90 Vol%, 96% Vol% (in distilled water), 100 Vol% twice at 35°C for 12 hours each step, followed by immersion in dichloromethane DCM (Sigma, 270997) for 45-60 minutes at room temperature and finally incubated with the refractive index matching solution BABB-D15 containing 15 parts BABB (benzyl alcohol + benzyl benzoate 1:2, Sigma, 24122 and W213802), 1 part diphenyl ether (DPE) (Alfa Aesar, A15791) and 0.4% Vol vitamin E (DL-alpha-tocopherol, Alfa Aesar, A17039), for at least 6 hours at room temperature until achieving transparency.

### vDISCO whole-body immunostaining, PI labeling and clearing

The following nanobodies and dyes were used for whole body immunostaining: Atto594 conjugated anti RFP nanobooster (Chromotek, rba594-100), Atto647N conjugated anti GFP nanobooster (Chromotek, gba647n-100), Atto488N conjugated anti GFP nanobooster (Chromotek, gba488-100), Propidium iodide (PI, Sigma, P4864).

In order to remove remaining blood and heme after PFA perfusion, and to decalcify the bones, the animals were subjected to perfusion with decolorization solution and decalcification solution before immunostaining. The decolorization solution was made with 25-30 Vol% dilution of CUBIC reagent 1^5^ in 0.1 M PBS. CUBIC reagent 1 was prepared with 25 wt% urea (Carl Roth, 3941.3), 25 wt% *N,N,Ń,Ń-*tetrakis (2-hydroxypropyl)ethylenediamine (Sigma, 122262) and 15 wt% Triton X-100 (AppliChem, A4975,1000) in 0.1 M PBS. The decalcification solution consisted of 10 wt/Vol% EDTA (Carl Roth, 1702922685) in 0.1 M PBS adjusting the pH to 8-9 with sodium hydroxide NaOH (Sigma, 71687).

The solutions for the immunolabeling pipeline were pumped inside the body of the animal by transcardial-circulatory perfusion exploiting the same entry point hole into the heart created during the PBS and PFA perfusion step (see above, perfusion and tissue preparation paragraph) and following the procedure already described in Pan *et al*., 2016. In brief, the mouse body was placed in a 300 ml glass chamber (Omnilab, 5163279) filled with 250-300 ml of appropriate solution, which covered the body completely. Next, the transcardial-circulatory system was established involving a peristaltic pump (ISMATEC, REGLO Digital MS-4/8 ISM 834; reference tubing, SC0266) keeping the pressure at 180-230 mmHg (50-60 rpm). One channel from the pump, made by a single reference tube, was set for circulation of the solution through the heart into the vasculature: one ending of the tube was connected to the tip of a syringe (cut from a 1 ml syringe-Braun, 9166017V) which held the perfusion needle (Leica, 39471024) and the other ending was immersed in the solution chamber where the animal was placed. The perfusion needle pumped the appropriate solution into the mouse body, and the other ending collected the solution exiting from the mouse body in order to recirculate the solution, pumping it back into the animal. To fix the needle tip in place and to ensure extensive perfusion, we put a drop of super glue (Pattex, PSK1C) at the level of the hole where the needle was inserted inside the heart. Using the setting explained above, after post-fixation and PBS washing, the mice were first perfused with 0.1 M PBS overnight at room temperature, then the animals were perfused with 250 ml of decolorization solution for 2 days at room temperature, exchanging with fresh decolorization solution every 6-12 hours until the solution turned from yellowish to clear and the spleen became lighter colour (indicating that the blood heme was extracted). Then, they were perfused with 0.1 M PBS, washing for 3 hours 3 times, followed by perfusion with 250 ml of decalcification solution for 2 days at room temperature and again perfused/washed with 0.1 M PBS for 3 hours 3 times. After this, the animals were perfused with 250 ml of permeabilization solution containing 1.5% goat serum (Gibco, 16210072), 0.5% Triton X-100, 0.5 mM of Methyl-beta-cyclodextrin (Sigma, 332615), 0.2% trans-1-Acetyl-4-hydroxy-L-proline (Sigma, 441562) and 0.05% Sodium Azide (Sigma, 71290) in 0.1 M PBS for half a day at room temperature. Subsequently, the perfusion proceeded further, connecting a 0.20 μm syringe filter (Sartorius, 16532) to the ending of the tube not holding the needle, in order to efficiently prevent accumulation of dye aggregates into the sample. At the same time, from this step we used an infrared lamp (Beuer, IL21) directed to the chamber to heat up the solution to 26-28°C. With this setting, the animals were perfused for 6 days with 250 ml of the same permeabilization solution containing 35 μl of nanobooster (stock concentration 0.5 – 1 mg/ml) (the amount of nanobody was adjusted depending on the expected presence of fluorescent protein in the mouse body) and 290 μl of propidium iodide (stock concentration 1 mg/ml). Next, we removed the animals from the chamber and with fine scissors we removed a tiny piece of skull from the back of the skull (above the cerebellum) at the level of the occipital bone, and we placed the bodies in a 50 ml tube (Falcon, 352070), filled with the same permeabilization solution, containing an extra 5 μl of nanobooster and incubated the tubes at 37°C with gentle shaking for an additional 2-3 days of labeling. After that, the mice were placed back in the perfusion system and labeling solution was washed out by perfusing with washing solution (1.5% goat serum, 0.5% Triton X-100, 0.05% of sodium azide in 0.1 M PBS) for 3 hours per 3 times at room temperature and 0.1 M PBS for 3 hours per 3 times at room temperature.

After the staining, the animals were cleared using a 3DISCO based passive whole-body clearing protocol optimized for big samples. The mice were incubated at room temperature in dehydration and clearing solutions inside a 300 ml glass chamber, kept with gentle rotation on top of a shaking rocker (IKA, 2D digital) inside a fume hood. For dehydration, mice bodies were incubated in 200 ml of the following gradient of tetrahydrofuran THF (Sigma, 186562) in distilled water (12 hours for each step): 50 Vol% THF, 70 Vol% THF, 80 Vol% THF, 100 Vol% THF and again 100 Vol% THF, followed by 3 hours in dichloromethane and finally in BABB. During all incubation steps, the glass chamber was sealed with parafilm and covered with aluminum foil.

### vDISCO whole-mount immunolabeling of individual organs

For the quantification of fluorescence signal after nanoboosting, dissected brains were stained using the immunolabeling protocol for dissected organs: first the post-fixed brains were pretreated, incubating them for 2 days at 37°C with gentle shaking in 4.5 ml of same solution used at the permeabilization step (see paragraph above) (1.5% goat serum, 0.5% Triton X-100, 0.5 mM of Methyl-beta-cyclodextrin, 0.2% trans-1-Acetyl-4-hydroxy-L-proline, 0.05% sodium azide in 0.1 M PBS). Subsequently, the brains were incubated in 4.5 ml of this same permeabilization solution + Atto647N conjugated anti GFP nanobooster (dilution 1:600) for 12-14 days at 37°C with gentle shaking, then brains were washed for 2 hours 3 times and once overnight with the washing solution (1.5% goat serum, 0.5% Triton X-100, 0.05% of sodium azide in 0.1 M PBS) at room temperature and in the end washed for 2 hours 4 times with 0.1 M PBS at room temperature. The immunostained brains were cleared with 3DISCO clearing: first they were put in the Eppendorf 5 ml tubes and then incubated at room temperature with gentle shaking in 4.5 ml of the following gradient of THF in distilled water (2 hours for each step): 50 Vol% THF, 70 Vol% THF, 80 Vol% THF, 100 Vol% THF and overnight 100 Vol% THF; after dehydration, the samples were incubated for 1 hour in dichloromethane, and finally in BABB until transparency. During all the clearing steps, the tubes were wrapped with aluminum foil to keep them in dark.

### Light-sheet microscopy imaging

Single plane illuminated (light-sheet) image stacks were acquired using an Ultramicroscope II (LaVision BioTec), featuring an axial resolution of 4 μm with following filter sets: ex 470/40 nm, em 535/50 nm; ex 545/25 nm, em 605/70 nm; ex 560/30 nm, em 609/54 nm; ex 580/25 nm, em 625/30 nm; ex 640/40 nm, em 690/50 nm. For low magnification-whole-body imaging of the *Thy1*-GFPM mouse, we used a 1x Olympus air objective (Olympus MV PLAPO 1x/0.25 NA [WD = 65mm]) coupled to an Olympus MVX10 zoom body, which provided zoom-out and -in ranging from 0.63x up to 6.3x. Using 1x objective and 0.63x of zoom, we imaged a field of view of 2 × 2.5 cm, covering the entire width of the mouse body. Tile scans with 60% overlap along the longitudinal y-axis of the mouse body were obtained from ventral and dorsal surfaces up to 13 mm in depth, covering the entire volume of the body using a z-step of 8 μm. Exposure time was 120 ms, laser power was adjusted depending on the intensity of the fluorescent signal (in order to never reach the saturation) and the light-sheet width was kept at maximum. After tile imaging of the sample within the entire field of view, already scanned regions were cut using a thin motorized dental blade (0.2 mm) (Dremel 8200) for further imaging. After low magnification imaging of the whole body, a forelimb of the *Thy1*-GFPM animal was imaged with a 2x objective (Olympus MVPLAPO2XC/0.5 NA [WD = 6 mm]) coupled with the same Olympus MVX10 zoom body at zoom magnification 1.6x. Moreover, the same 2x objective was used to perform high magnification imaging of specific body regions (e.g back of the animal at the level of lumbar vertebra or the head). Individual organs (including brain, lungs, intestine and thymus) were imaged individually using high magnification objectives: 2x objective (Olympus MVPLAPO2XC/0.5 NA [WD = 6 mm]) coupled with the same Olympus MVX10 zoom body, 4x objective (Olympus XLFLUOR 4x corrected/0.28 NA [WD = 10 mm]), 25x objective (Olympus XLPLN 25x/0.95 NA [WD 4mm]) and 20x objective (Zeiss 20x Clr Plan-Neofluar/0.1 NA [WD 4 = mm]) coupled to an Olympus revolving zoom body unit (U-TVCAC) kept at 1x. High magnification tile scans were acquired using 10-30% overlap and the light-sheet width was reduced to obtain maximum illumination in the field.

### Reconstructions of whole-mouse body scans

We acquired light-sheet microscope stacks using ImSpector (Version 5.295, LaVision BioTec GmbH) as 16-bit grayscale TIFF images for each channel separately. In each pair of neighbouring stacks, alignment was done by manually selecting 3 to 4 anatomic landmarks from the overlapping regions, then the stitching was done sequentially with the Scope Fusion module of the Vision4D (Version 2.12.6 x64, Arivis AG) software. Landmarks were mainly chosen from the skeletal bones or fewer from the neuronal structures based on visual inspection of the anatomical features. After completing the 3D reconstructions, the data visualization was done with Amira (Version 6.3.0, FEI Visualization Sciences Group), Imaris (Version 9.1, Bitplane AG) and Vision4D in both volumetric and maximum intensity projection color mapping. Depth coding was done using Temporal-Color Code plugin in Fiji.

### Neuron tracing

For automated neuron tracing in our light-sheet datasets obtained with Zeiss 20x Clr Plan-Neofluar/0.1 NA [WD 4 = mm] objective, we used NeuroGPS-Tree algorithm^24^. The NeuroGPS-Tree was developed for tracing relatively small volumes of confocal microscopy data, therefore, we initially reduced the file size to under 1 GB (approximate maximum data size for NeuroGPS-Tree computation) by using Fiji scale function. Due to high signal intensity discrepancy between soma and neurites, we next processed the data with a custom-made macro in Fiji (available upon request), which consisted of background removal, pseudo background correction, noise filtering and sharpening. Next, the pre-processed data was loaded and analysed first in NeuroGPS for soma detection and later in NeuroGPS-Tree for neurite detection (both steps are part of the same algorithm package). The best parameters of soma and neurite detection were chosen following the original publication. To quantify the features (such as number and total length of neurites per cell) of these detected neuronal cells, we used Amira software: we chose the 10 neurons with the biggest file size per each group and we analyzed them using the Spatial Graph Statistics function of the software.

### Quantifications

#### Analysis of fluorescence signal profiles from light-sheet images

The fluorescence signal profiles from each channel (excitation 470 nm, 560 nm and 647 nm) were plotted in the same z-stack and normalized as percentage over the maximum peak using Fiji. (Supplementary Fig.1).

To compare vDISCO boosted vs. unboosted protocols and consequently the reduction of the background and the improvement of the signal over background ratio in far-red and near far-red channels, we analyzed neurons and axonal bundles expressing EGFP imaged with excitation at 470 nm, and neurons and axonal bundles labelled with anti-EGFP nanobody conjugated with Atto647N imaged with excitation at 640 nm at the same anatomic region (n=9 neuronal structures per each experimental group which consisted of 3 animals per each imaging modality). The signal profile was analyzed in Fiji and measured from a defined straight line covering the neuronal structure and surrounding tissue background and the normalized plots of the signal profile (Fig. 1k) were calculated by normalizing the plots of neuronal structures obtained as described above over the average signal intensity of the respective surrounding background. Each experimental group consisted of 3 animals and per each animal at the same anatomic region we plotted 3 profiles.

#### Fluorescence level

Fluorescence level quantification was expressed as a signal-to-background ratio and was calculated using Fiji^19^ at the following time points: 0, 2, 3, 4 and half, 5 and half, 12 and 18 months after nanoboosting (Fig. 1p,q). Each 4x light-sheet microscopy brain scan was taken with the same imaging parameters, and an image in Tiff format of the same anatomic region for all the samples was quantified. The mean value of the background for each image was obtained by averaging the background values of 12-40 regions from equally sized areas of the image in regions of the sample without signal. To calculate the mean value of the signal per each image, we used the threshold function of the software: the threshold was adjusted to consider the fluorescence signal visible in the image. After adjusting the threshold, only the sharp signal from specific cellular structures was analyzed per image. To this end, we used Fiji’s “analyse particles” function to measure the signal intensity only of particles sized between 5-10 and 100-150 pixels (visible fluorescent cells), and calculated the average value from all the particles. Next, this value was divided by the mean value of the background of the respective image, obtaining the fluorescence level over the background. The corresponding images, visually showing the preservation of signal over time in relation to the respective fluorescence levels, were processed using Enhance Local Contrast (CLAHE) function in Fiji to increase the contrast of fluorescent cells in the tissue.

#### ClearMap

To quantify microglia distribution we used ClearMap^26^. Since the script was originally developed for quantification of the cFos+ cells, to comply with offered method, we did the following pre-processing steps on our microglia data using Fiji before ClearMap:

- Background equalization to homogenize intensity distribution and appearance of the microglia cells over different regions of the brain, using pseudo flat-field correction function from Bio-Voxxel toolbox.
- Convoluted background removal, to remove all particles bigger than relevant cells. This was done with the median option in Bio-Voxxel toolbox.
- 2D median filter to remove remaining noise after background removal. The filter radius was chosen to ensure the removal of all particles smaller than microglia cells.
- Unshapen mask to amplify the high-frequency components of a signal and increase overall accuracy of the cell detection algorithm of ClearMap.

After pre-processing, ClearMap was applied by following the original publication and considering the threshold levels that we obtained from the pre-processing steps. As soon as the quantification was completed, the data was exported as excel file for further analysis. For example, the cellular density per each brain region was obtained considering the absolute number of cells detected by ClearMap and the volume of that specific brain region, which was calculated using a custom script (available upon request) based on ClearMap (Elastix registration).

#### Quantification of peripheral neuronal degeneration in acute brain injury

Peripheral neuronal degeneration in TBI animals versus unlesioned control animals was assessed considering the complexity of axonal ramifications that projected from the left cervical and thoracic vertebra to the left muscles of the back at the level of the torso of the mouse, including the left spinotrapezius and latissimus dorsi. The complexity was expressed as number of axonal endpoints (nerve terminals that appear as button-like shape (See Fig. 3) over the total length of axonal ramifications that were protruding from a main branch. To calculate this index, first a 3×3 tile z-stacks of this anatomic region was taken from the animals by light-sheet microscopy using the 2x objective described in “light-sheet microscopy imaging section” (Olympus MVPLAPO2XC/0.5 NA objective coupled with the Olympus MVX10 zoom body) with a total magnification of 5x, in order to have enough resolution to manually trace the axonal ramifications and axonal end-feet. Then, the analysis was done over the max intensity projections of the tile scans with Fiji software. To measure the length of the ramifications, we used the “free hand line” function and the “ROI manager tool” of Fiji, in order to record all the traced axonal ramifications of interest, which were coming from a main branch; later we calculated the sum of the length of all of the recorded ramifications with “measure” function of Fiji. To count the nerve terminals, we used the “point tool” function and the same “ROI manager tool” in order to record all the visible axonal endpoints protruding from the traced ramifications. The analysis was performed over 2-4 branches from this same anatomic region per each animal, considering 3 animals per each experimental group.

### Statistical analysis

Data are presented as mean ± s.d. The data distribution in each experiment was checked for normality using Shapiro-Wilk Test. *P* values were calculated using two tailed unpaired *t*-test to compare data between two groups. A *p* value <0.05 was considered statistically significant.

## VIDEO LEGENDS

### Video 1

3D reconstruction of neuronal connectivity in a *Thy1*-GFPM mouse scanned by light-sheet microscopy. The muscles (red) are visualized in autofluorescent channel (blue-green spectra). The bones and internal organs (white) are prominent with PI labeling compared to other tissues in the body. The EGFP expressing neurons are boosted with nanobody conjugated with Atto 647N dye and imaged in far-red channel. The overall view of entire labeled nervous system in *Thy1*-GFPM mouse and fine details of neuronal connections are evident throughout the whole body.

### Video 2

3D visualization of neuronal connections from spinal cord to right forelimb obtained by light-sheet microscopy in *Thy1*-GFPM mouse. The muscles are shown in red, bones in white and the neurons in green. The fine details of axonal extensions and their endings at neuromuscular junctions are visible.

### Video 3

The first part of the video is the 2D orthoslicing of spinal cord of an intact *Thy*1-GFPM mouse in dorsoventral orientation. The muscles are shown in red, bones in white and the neurons in green. The details of neuronal cell bodies in ganglia embedded in the spinal cord vertebra and their axonal extensions into the CNS and PNS are visible. In the second part, neuronal connections (green) from spinal cord to muscles are shown in 3D and 2D.

### Video 4

Multiplexed visualization of CX3CR1^GFP/+^ x CCR2^RFP/+^ transgenic mouse head after panoptic imaging by two different nanoboosters (anti-GFP conjugated to Atto-647N and anti-RFP conjugated to Atto-594N). The CX3CR1 GFP+ microglia cells in the brain parenchyma vs. CCR2 RFP+ peripheral immune cells in the meningeal vessels were clearly visible in 3D reconstruction and 2D orthoslicing.

### Video 5

Panoptic imaging of *CD68*-EGFP transgenic mice upon SCI. 2D orthoslicing in horizontal and sagittal views clearly shows the activated immune cells in the muscles, spinal cord roots and meningeal compartments.

### Video 6

A *Thy1*-GFPM mouse brain was imaged by light-sheet microscopy more than a year after vDISCO. The details of the neuronal structures down to subcellular level were imaged by the Zeiss 20x on the light-sheet microscope.

---

**Supplementary Figure 1.**
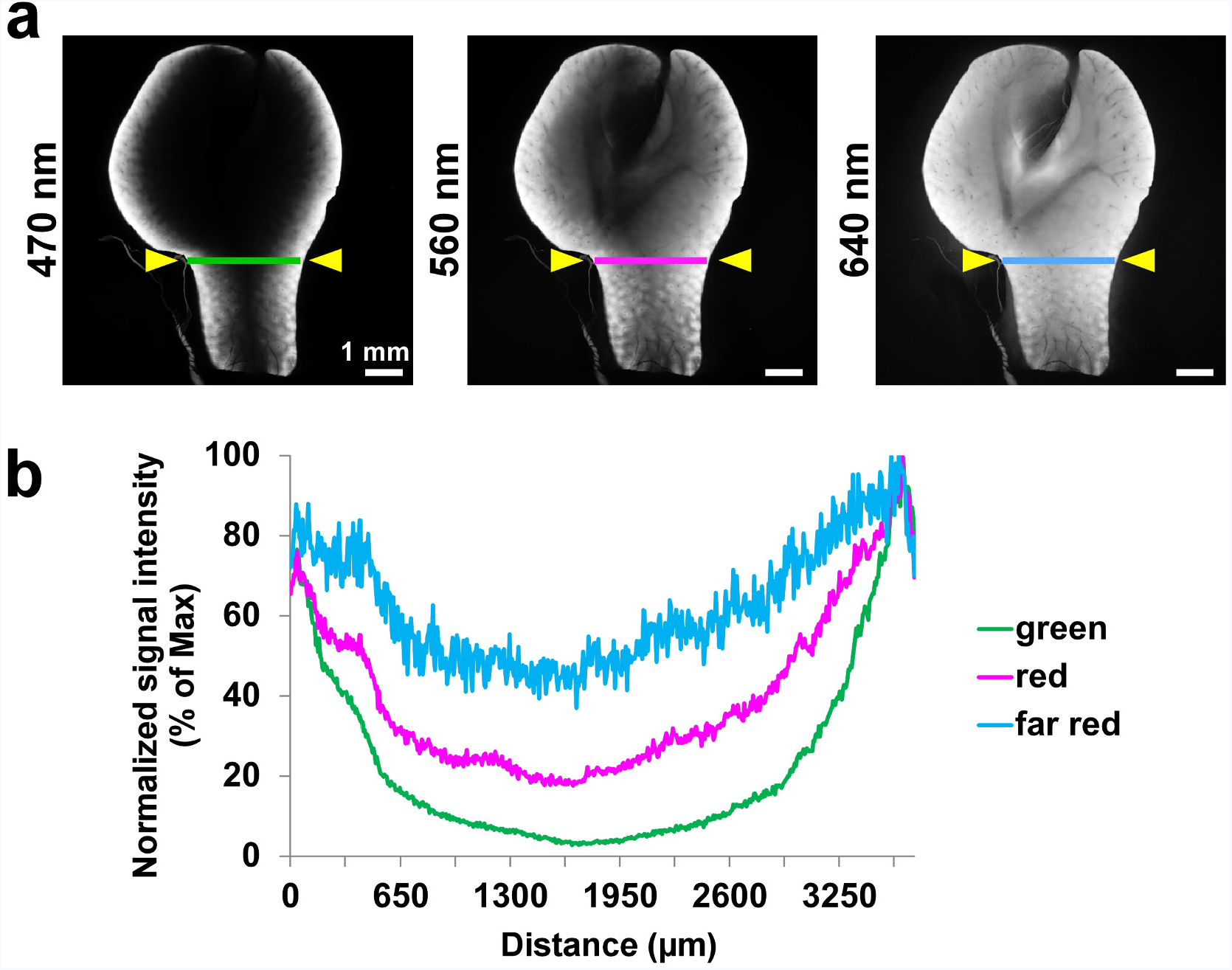
Light penetration deep in tissue at different wavelengths. (**a**) Demonstration of tissue penetration of the light at different imaging wavelengths. The same liver region (without any labeling) of a cleared mouse imaged in green (ex: 470 nm) (left), red (ex: 560 nm) (middle) and far-red channels (ex: 640 nm) (right) using light-sheet microscopy. (**b**) Fluorescence signal intensity profiles normalized over the maximum intensity of the regions indicated by the lines in **a**. Complete illumination of cleared liver at 640 nm compared to other wavelengths is evident.

**Supplementary Figure 2.**
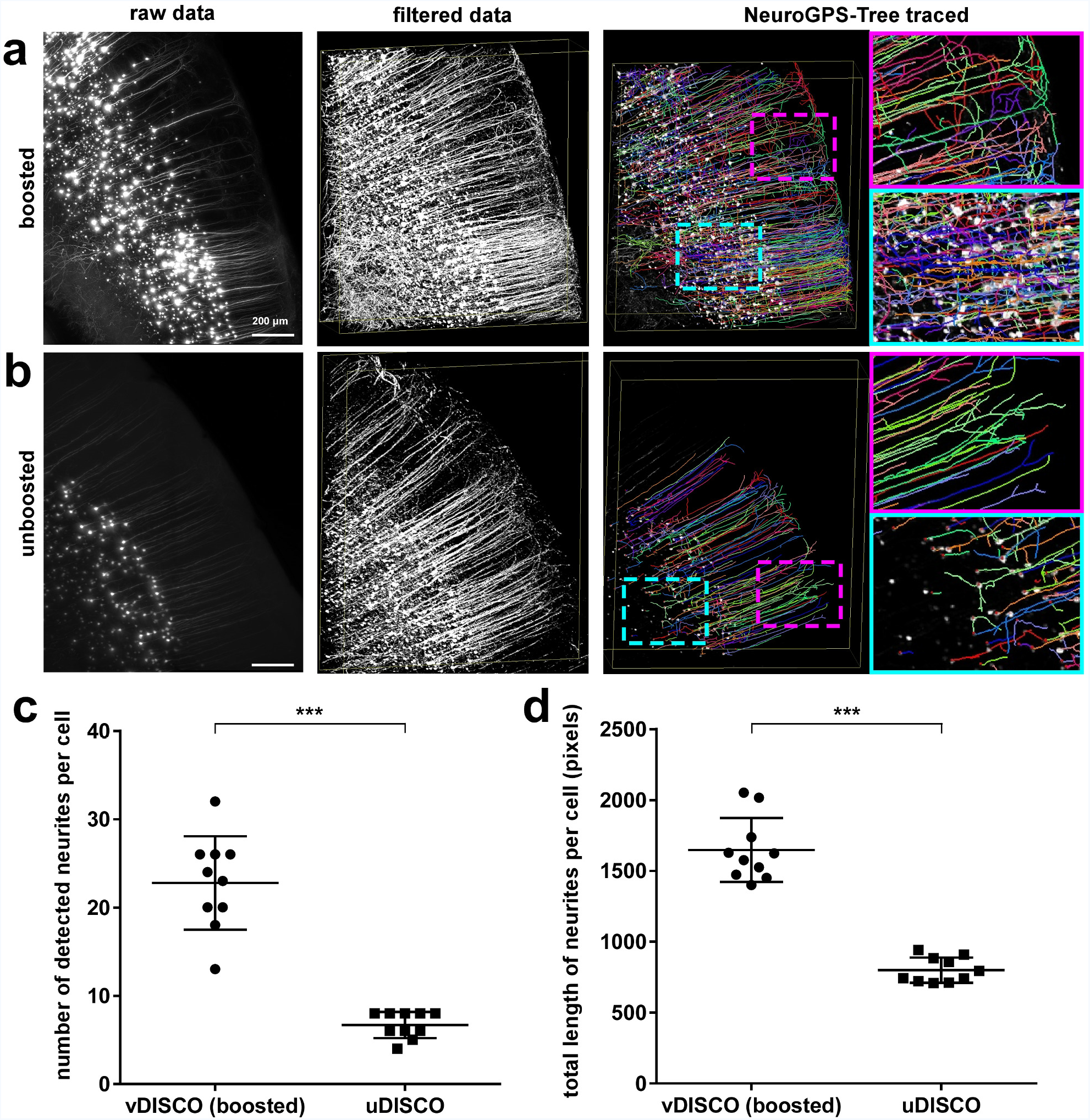
Neuron tracing with NeuroGPS-Tree. (**a,b**) Tracing of neurons in vDISCO boosted (**a**) and unboosted (**b**) brains from Thy1-GFPM mice using NeuroGPS-Tree algorithms. The scans were obtained with a commercial light-sheet microscope (Ultramicroscope II, LaVision Biotec), optimized for large cleared tissue imaging, thereby with lower resolution compared to standard confocal microscopes. From left to right: raw image, equalized & filtered images, and NeuroGPS-Tree traced neuronal structures are shown: all the neurites belonging to a neuron are represented by the same color. Neuronal soma identified by NeuroGPS-Tree are represented by red dots. (**c**) Quantification of number of detected neurites per cell in vDISCO boosted vs. unboosted samples. (**d**) Quantification of total length of neurites per cell in vDISCO boosted vs. unboosted samples. (mean ± s.d; n=10 neurons per group; statistical significance (***p < 0.001) was assessed by two tailed t-test).

**Supplementary Figure 3.**
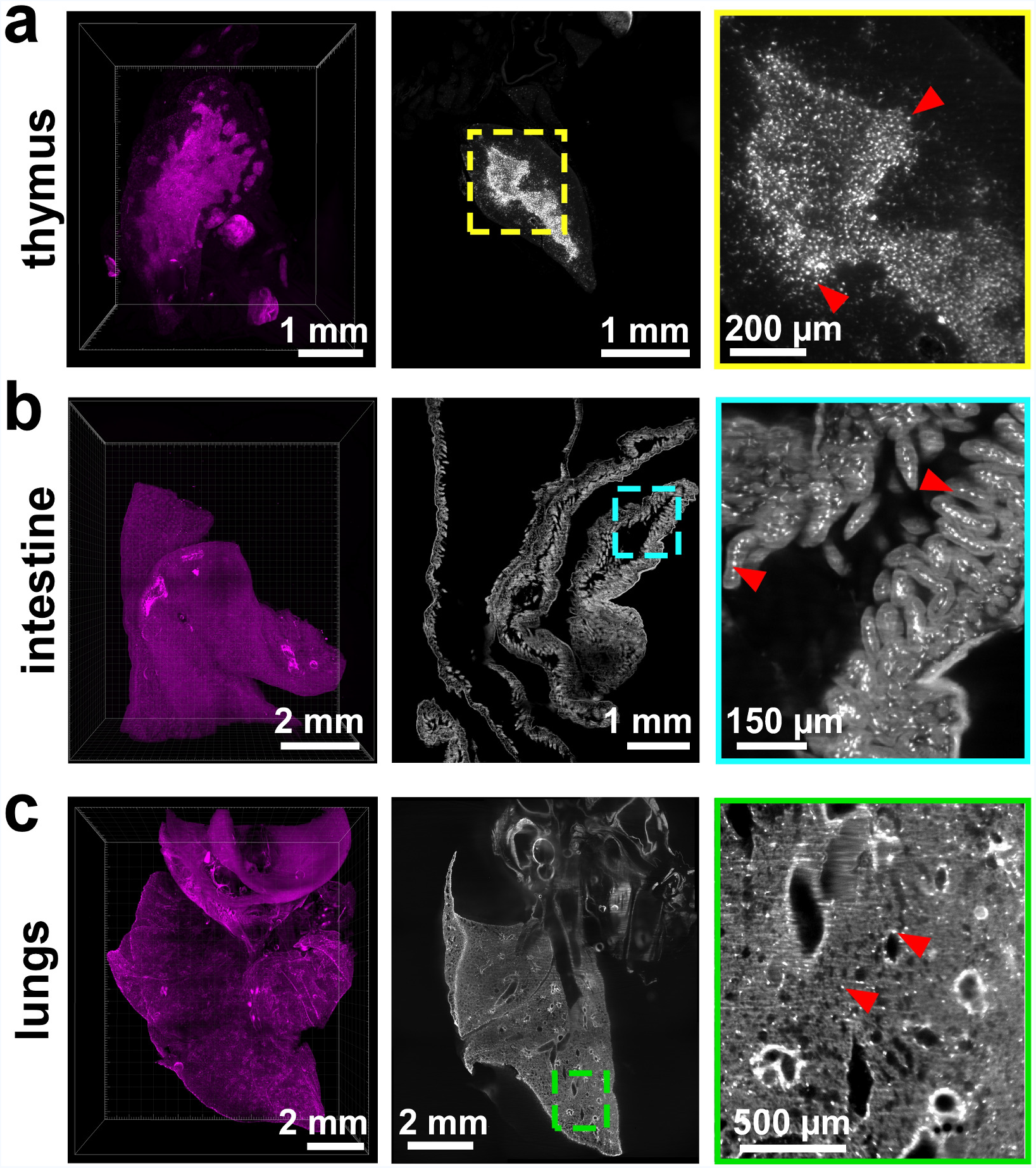
Visualization of CX3CR1 GFP+ cells in individual organs. (**a-c**) 3D visualization (magenta) and raw data images (grey) of individual organs including thymus (**a**), intestine (**b**), and lungs (**c**) imaged by light-sheet microscopy. Individual CX3CR1 GFP+ cells are indicated by red arrowheads in the zoomed images from the boxed regions.

**Supplementary Figure 4.**
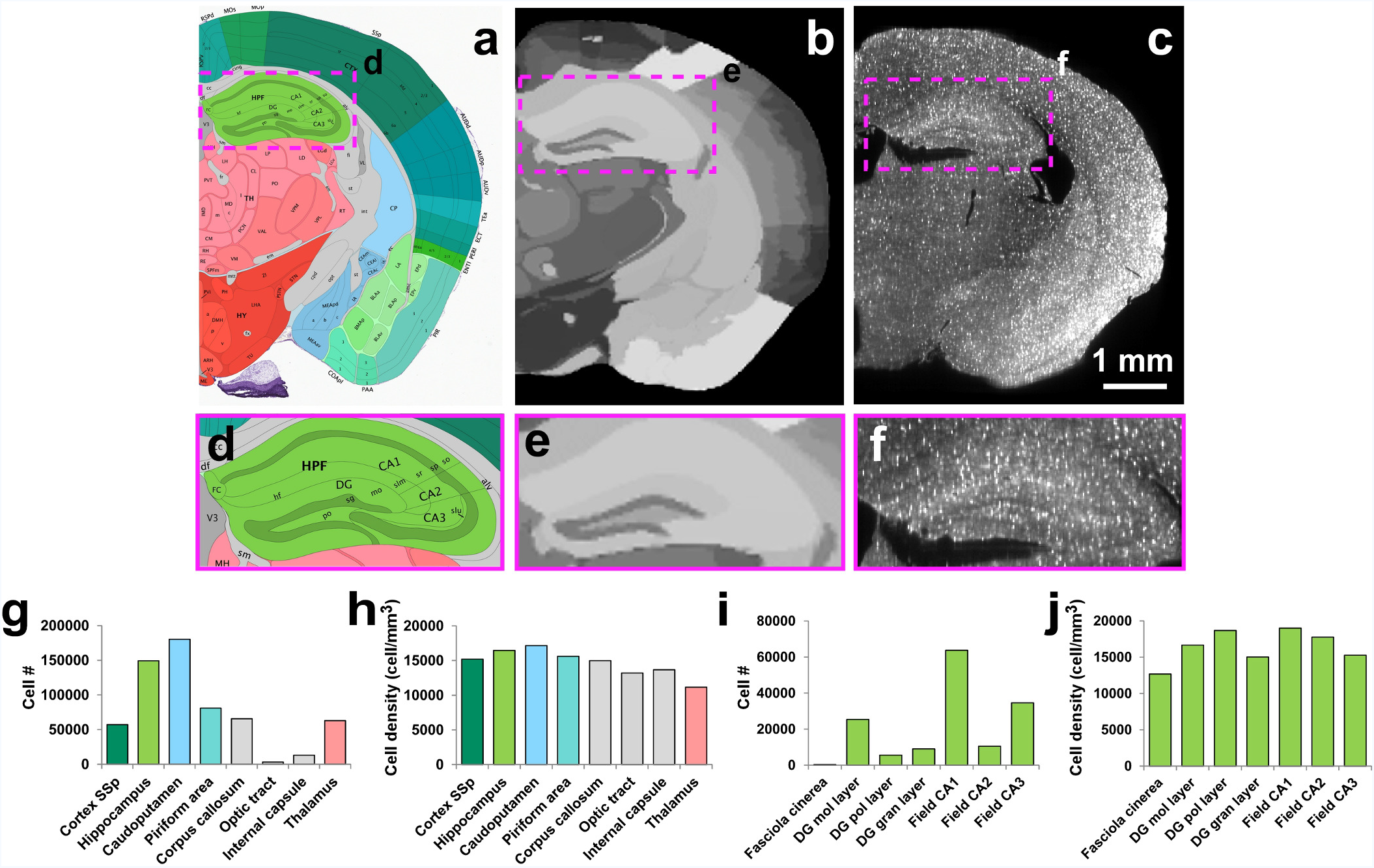
Visualization and quantification of the signal from CX3CR1GFP/+ mouse by vDISCO boosting. (**a-j**) Automated analysis of fluorescent cells distribution in CX3CR1GFP/+ mouse brain after vDISCO boosting (n = 2 mice, total number of microglia: 2.64 and 2.55 million, respectively): (**a**) reference image from the Allen brain atlas, (**b**) registered reference annotation image, (**c**) the corresponding region in raw data. (**d-f**) High magnification images of the region indicated in a-c containing the hippocampus. Automated quantification of absolute cell numbers (**g**) and cell densities (**h**) of the anatomical regions visible in a-c. Absolute cell numbers (**i**) and cell densities (**j**) of sub-regions of hippocampus showed in d-f. The quantifications include the cells in both brain hemispheres.

**Supplementary Figure 5.**
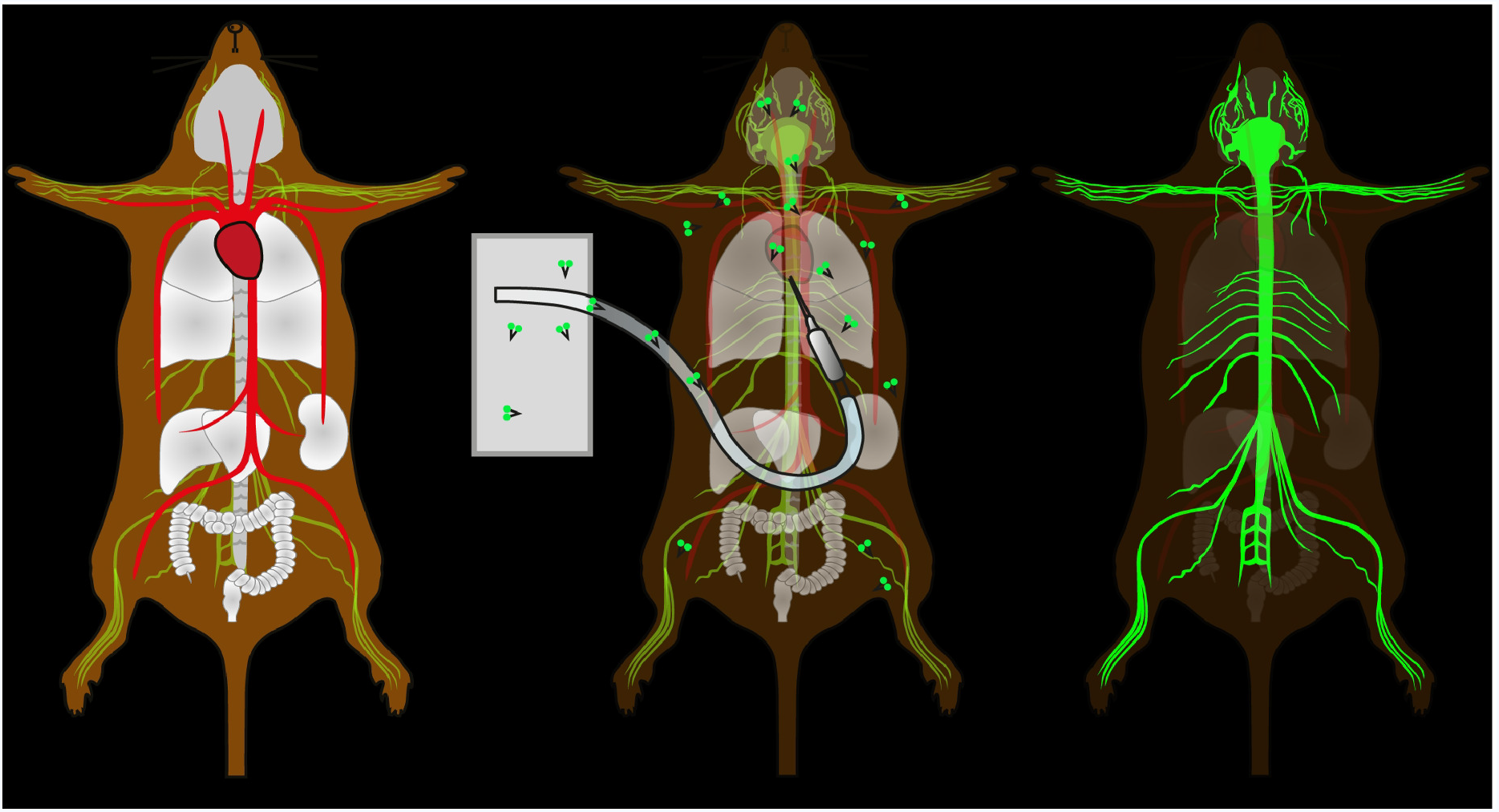
Schematic illustration of whole-body vDISCO boosting method. The decolorization, decalcification, and nanoboosting steps are performed via transcardiac perfusion. After boosting, the boosted fluorescence signal becomes highly visible over the reduced background in the intact transparent animals.

**Supplementary Figure 6.**
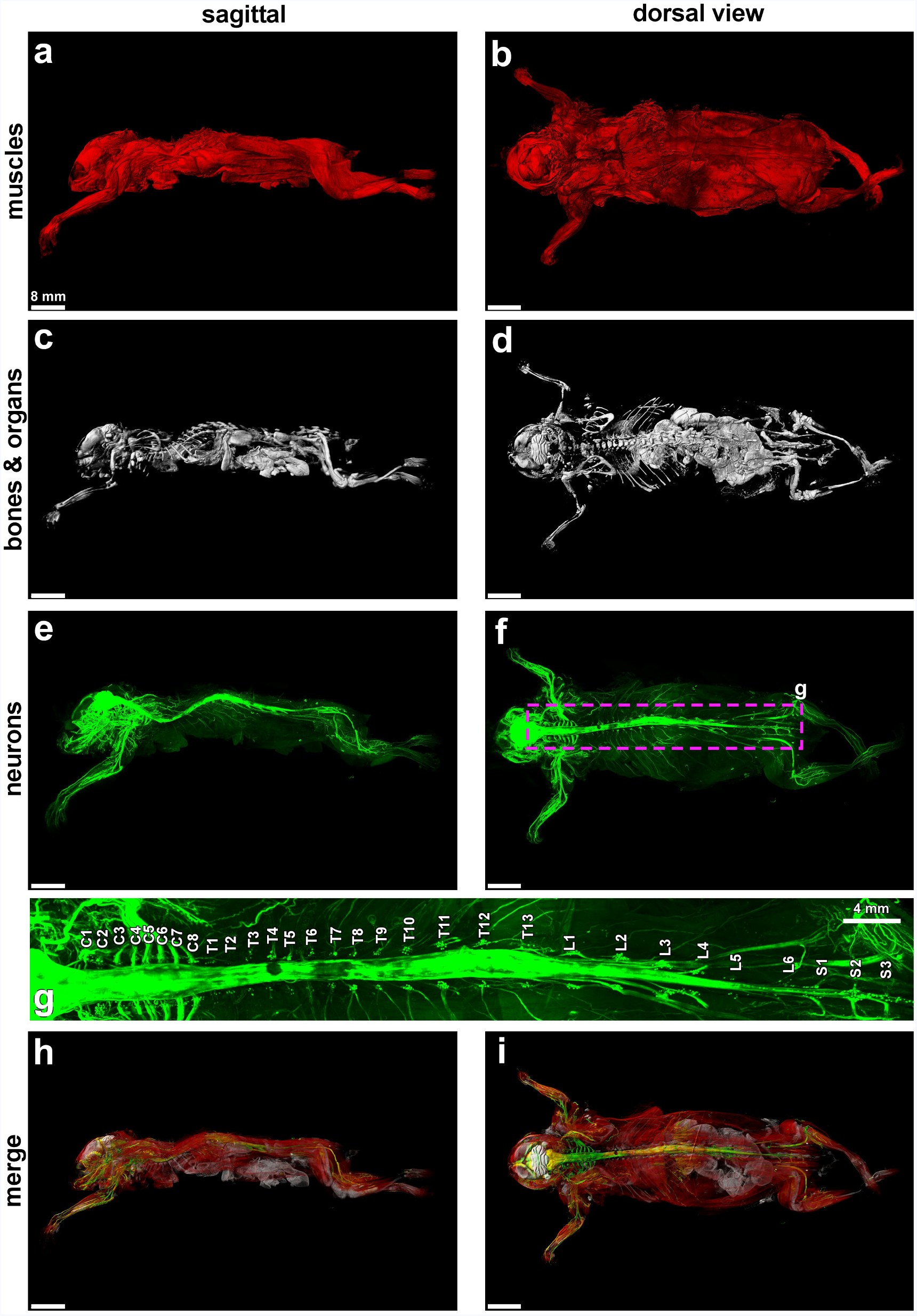
Whole body neuronal connectivity of a Thy1-GFPM mouse by panoptic imaging. (**a-d**) The sagittal and horizontal views of autoflourescence (muscles), and PI (bones and organs) channels. (**e,f**) Neuronal connectivity of the whole body in sagittal and horizontal views. (**g**) High magnification view from the indicated region in f showing details of spinal cord segments. (**h,i**) The merge channels in sagittal (**h**) and horizontal (**i**) views of neurons, bones and organs, and muscles are shown.

**Supplementary Figure 7.**
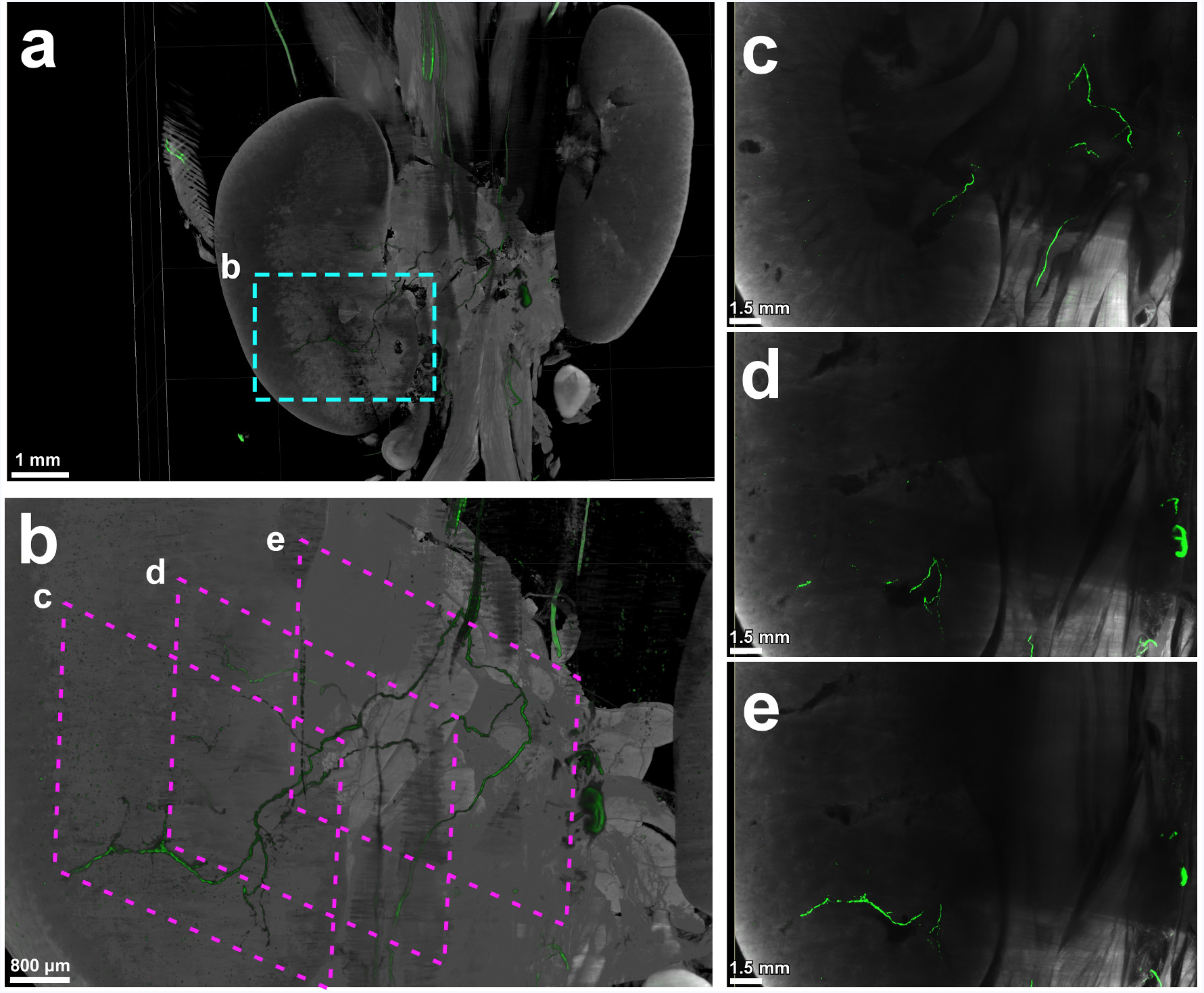
Peripheral nerves innervating kidneys in Thy1-YFPH mouse detected by panoptic imaging. (**a**) 3D visualization of nerves innervating the kidneys of a Thy1-YFPH animal. (**b**) Zoom image from the marked region in a. (**c-e**) 2D projection images of the kidney at the indicated depths in b.

**Supplementary Figure 8.**
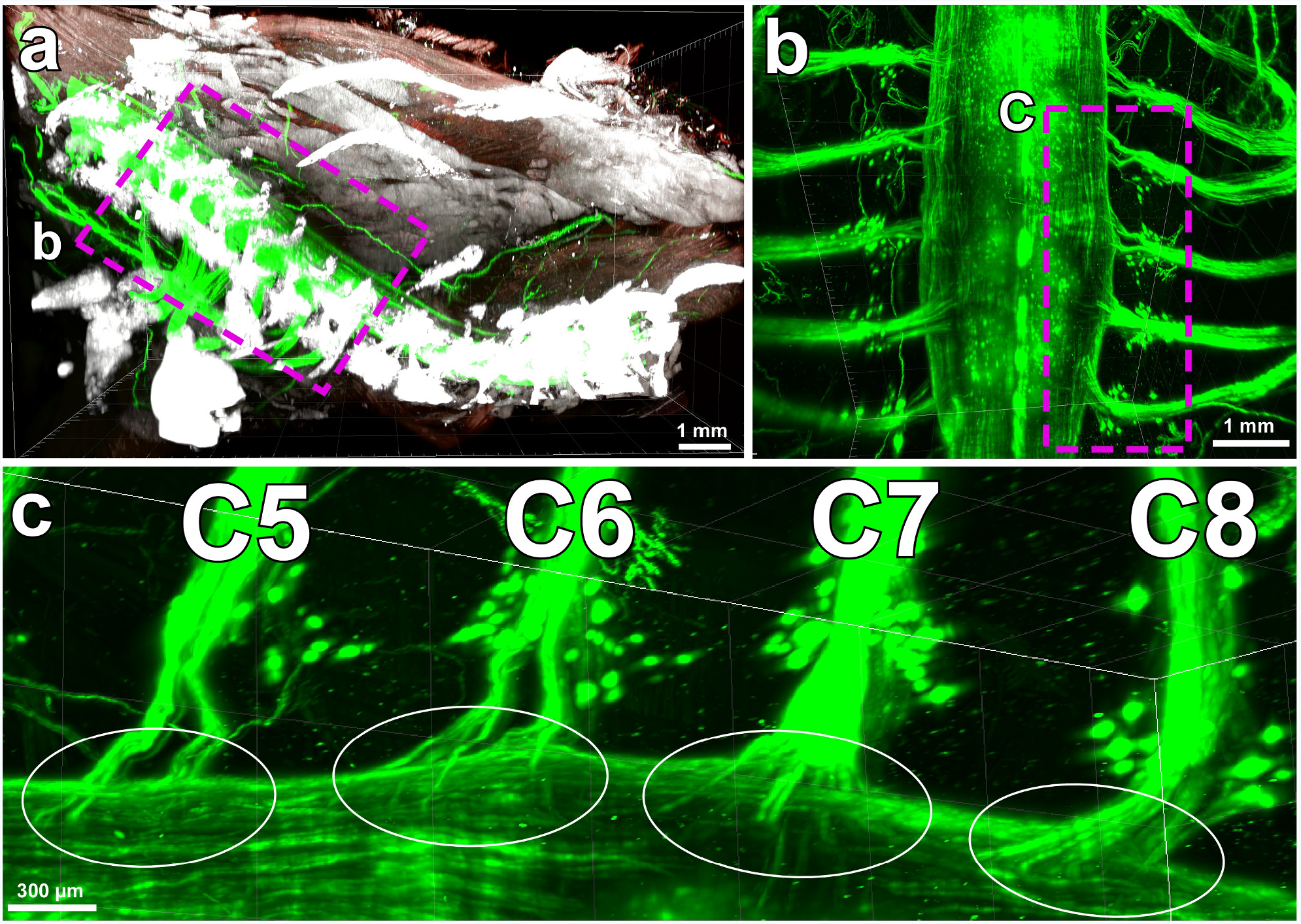
Upper torso and individual spinal cord roots. (**a**) Panoptic imaging of the upper torso of the transparent mouse showing individual spinal cord roots. The bones and organs are in white, the muscles are in red, and the nerves are in green. (**b**) High magnification view from the region indicated in a showing only nerve signal (ventral roots). (**c**) High magnification view from the region indicated in b demonstrating non-overlapping entry of axons in C5-C8 ventral spinal cord segments (white circles).

**Supplementary Figure 9.**
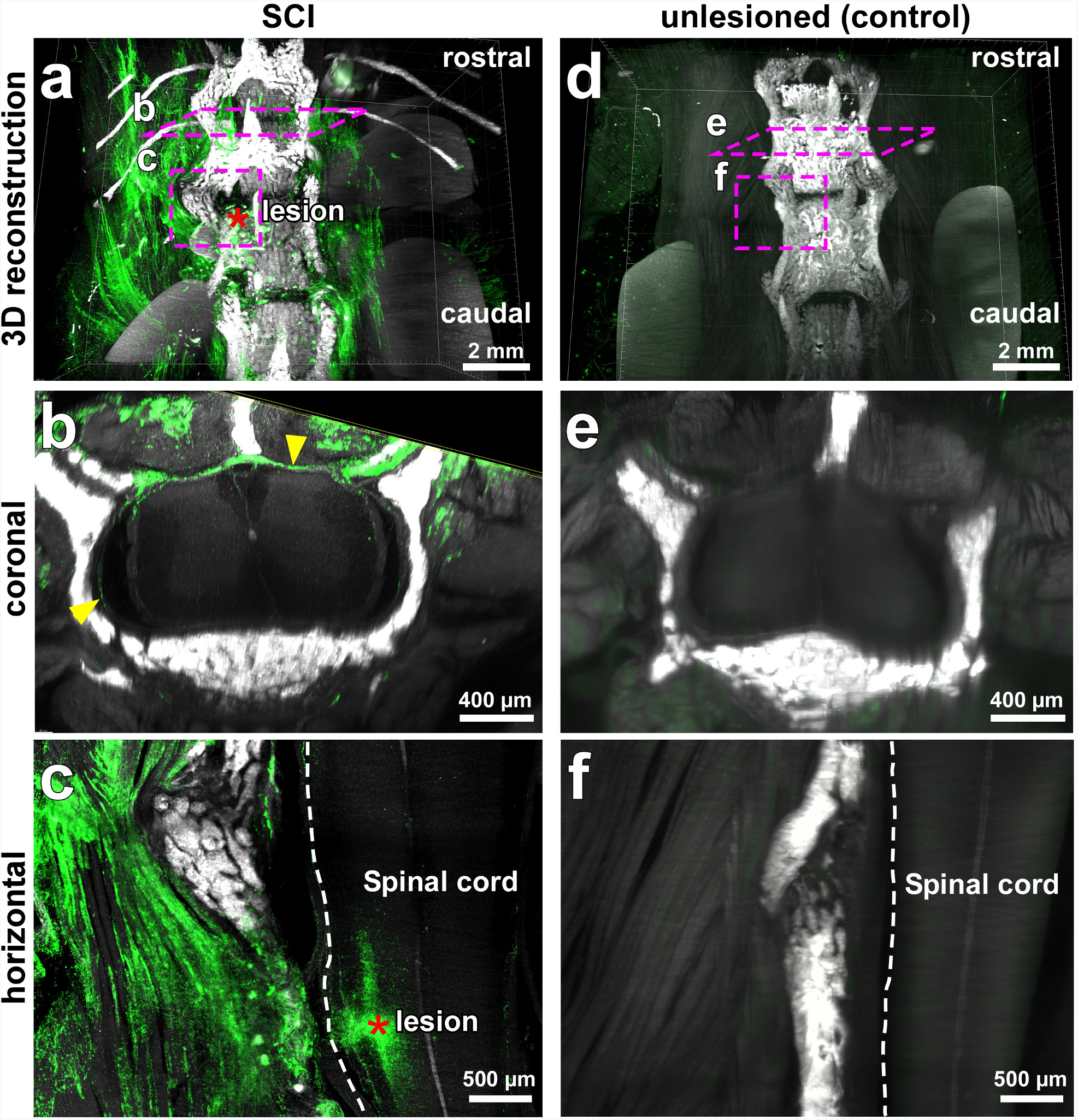
Immune cell activation and invasion induced by spinal cord injury. (**a-f**) 3D visualization of spinal cord from CD68EGFP/+ transgenic mice with spinal cord injury compared to unlesioned controls. CD68 GFP+ cells are shown in green and PI labeled bones in white. Red asterisks indicate the lesion site. Increased CD68 GFP+ cells throughout the muscles, spinal cord roots and meninges (yellow arrowheads) are evident in the injured spinal cord (**a-c**) compared to the controls (**d-f**). See also Video 5.

